# Reconstructing cell histories in space with image-readable base editor recording

**DOI:** 10.1101/2024.01.03.573434

**Authors:** Duncan M Chadly, Kirsten L Frieda, Chen Gui, Leslie Klock, Martin Tran, Margaret Y Sui, Yodai Takei, Remco Bouckaert, Carlos Lois, Long Cai, Michael B. Elowitz

**Affiliations:** Division of Biology and Biological Engineering, California Institute of Technology, Pasadena, CA, USA; Spatial Genomics, Pasadena, CA, USA; Palo Alto Veterans Institute for Research, Palo Alto, CA, USA; School of Computer Science, University of Auckland, New Zealand; Howard Hughes Medical Institute and Department of Applied Physics, California Institute of Technology, Pasadena, CA, USA

## Abstract

Knowing the ancestral states and lineage relationships of individual cells could unravel the dynamic programs underlying development. Engineering cells to actively record information within their own genomic DNA could reveal these histories, but existing recording systems have limited information capacity or disrupt spatial context. Here, we introduce *baseMEMOIR*, which combines base editing, sequential hybridization imaging, and Bayesian inference to allow reconstruction of high-resolution cell lineage trees and cell state dynamics while preserving spatial organization. BaseMEMOIR stochastically and irreversibly edits engineered dinucleotides to one of three alternative image-readable states. By genomically integrating arrays of editable dinucleotides, we constructed an embryonic stem cell line with 792 bits of recordable, image-readable memory, a 50-fold increase over the state of the art. Simulations showed that this memory size was sufficient for accurate reconstruction of deep lineage trees. Experimentally, baseMEMOIR allowed precise reconstruction of lineage trees 6 or more generations deep in embryonic stem cell colonies. Further, it also allowed inference of ancestral cell states and their quantitative cell state transition rates, all from endpoint images. baseMEMOIR thus provides a scalable framework for reconstructing single cell histories in spatially organized multicellular systems.

## Introduction

Cells divide, differentiate, and move to form exquisitely organized structures. Reconstructing the dynamic histories of individual cells, particularly their lineage relationships, could enable researchers to understand how tissues form, analyze the roles of intrinsic and extrinsic determinants of cell fate decisions, and reveal how processes are dysregulated in disease^1^. Recent advances in single cell sequencing and spatial genomics now allow us to capture single cell states at specific moments in time^2–4^. However, with a few exceptions^5^, the histories of those cells have largely remained hidden.

Researchers have sought to address this challenge through engineered recording systems, which progressively introduce stochastic edits in genomically integrated barcode sequences as cells proliferate. Systems such as GESTALT^6–8^, CARLIN^9^, LINNAEUS^10^, SMALT^11^, and the homing CRISPR barcoded mouse^12,13^, use CRISPR/Cas9 or recombinases to edit designed target sequences, relying on next generation sequencing to read out edited barcodes.

Alternative systems, including CAMERA, leverage CRISPR base editors to generate more specific types of barcode diversity.^14,15^ Further, prime editors introduced an additional paradigm for phylogenetic recording, sequentially inserting short nucleotide motifs for genomic information storage.^16–18^ In all of these systems, lineage relationships between individual cells are reconstructed from each cell’s unique pattern of target site edits, in a manner analogous to sequence-based phylogenetic reconstruction.^19,20^

These techniques can be powerful in their ability to recover lineage but disrupt spatial organization. A parallel set of methods were developed to allow barcode editing in ways that allow readout of edits and cell states by imaging^21,22^. For example, previous MEMOIR (Memory by Engineered Mutagenesis with Optical In situ Readout) systems showed that it is possible to stochastically and irreversibly edit engineered DNA barcodes, or ‘scratchpads,’ using CRISPR/Cas9^21^ or an integrase^22^, and then read out those edits by imaging. However, these methods were limited in accuracy by relatively low numbers of mutable target sites, which serve as memory in the genome. For example, existing image-readable systems have demonstrated only ∼16 bits of information storage^22^.

Recently, we showed that in situ T7 transcription can amplify genomic DNA into localized RNA clusters, which can then be competitively probed to discriminate single base edits^23^. In this strategy, termed “Zombie,” a genomic DNA of interest can be maintained without transcription in live cells, transcribed after fixation with the addition of T7 polymerase, and finally detected by RNA-FISH. Zombie transcription avoids silencing problems that occur when barcodes must be continuously transcribed in live cells, generates large quantities of RNA that spatially localize around the active site of transcription, can detect mutations at single nucleotide resolution, and is compatible with subsequent sequential rounds of FISH to detect endogenous gene transcripts. Zombie enables readout of dense editable memory arrays, expanding the capacity of MEMOIR approaches.

Here, we introduce “baseMEMOIR”, a multiplexed phylogenetic recording system which enables detailed recording of lineage relationships over time in a manner compatible with recovery of spatial position and gene expression patterns (**Figure 1a**). We distributed mutable synthetic DNA sequences at high copy number randomly across the genome of mouse embryonic stem cells. These targets can be edited by the CRISPR A-to-G base editor (ABE), which complexes with guide RNAs to specifically mutate target sites within our synthetic sequences. For tight control of editing, we used two inducible systems to control the base editor (the TRE3G Tet-On system) and guide RNAs (controlled by a Wnt responsive promoter) respectively. On induction, mutations occur at a rate commensurate with cell division and are passed down from parent cells to their progeny through DNA replication, creating lasting marks that link related cells to one another. We then recovered mutation states through microscopic imaging, and applied Bayesian phylogenetic tools to infer lineage relationships as well as transcriptional state dynamics and spatiotemporal histories in a unified manner. By comparing the distinct pattern of mutations in each cell after a series of divisions, we were able to reconstruct phylogenies and estimate uncertainty in both tree topology and the timing of past cell divisions.

**Figure 1:**
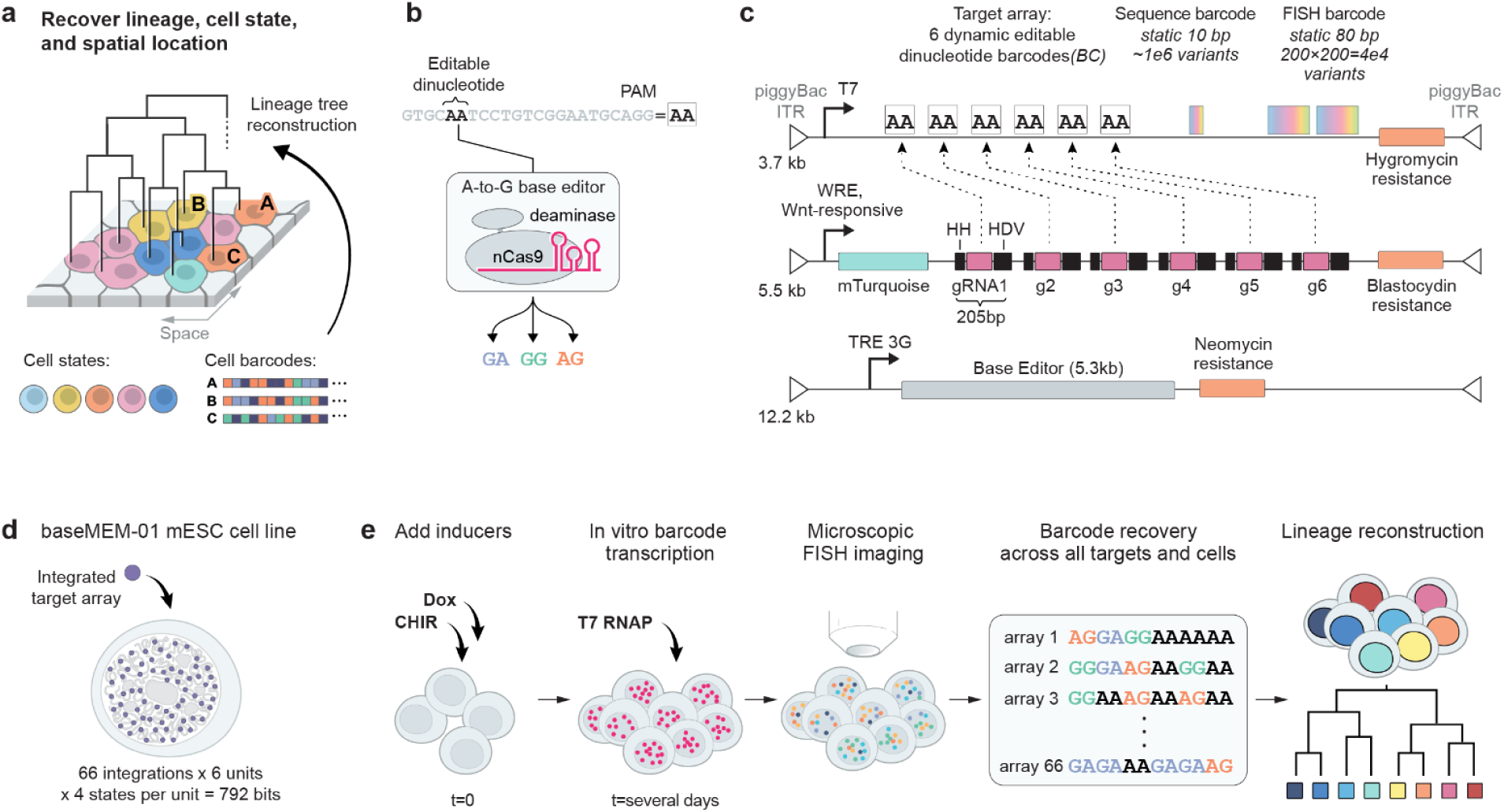
Multiplexed, genomically dispersed, editable barcodes enable detailed recording of lineages over many generations with in situ readout. **(a)** Detailed lineage trees can be measured alongside transcriptional cell states while maintaining spatial context through phylogenetic barcoding. **(b)** Predicted stochastic editing of AA dinucleotides results in one of three terminal outcomes. **(c)** An inducible barcode editing system can be integrated into cells at high copy number via piggyBac transposase. Target arrays (top) contain 6 AA dinucleotides flanked by unique protospacer sequences as well as sequencing and imaging-readable static barcodes which serve to uniquely mark different genomic integrations of the array. Editing is induced by expression of guide RNAs (middle), controlled by a Wnt-responsive element, and base editor (bottom), controlled by the TRE3G tet-on promoter. **(d)** We engineered a monoclonal mESC cell line containing 66 uniquely labeled target array copies (396 editable dinucleotides, or 792 bits of information) alongside the inducible editing machinery. **(e)** This cell line enables genomic lineage recording and recovery through FISH imaging and phylogenetic tree inference.

To demonstrate the capabilities of baseMEMOIR, we applied this system to estimate state switching rates and probable past cell states along lineages of dividing mESCs grown in serum-LIF conditions in the presence of a Wnt agonist. We find that mESCs grown in these conditions undergo reversible transitions between formative and 2C-like states, with an intermediate naive state that can be broken up into three distinct subclusters. Each state and subcluster, in addition to being transcriptionally distinct, is further distinguished by a set of allowed state transitions.

The baseMEMOIR cell lines and platform can be applied, and further scaled, in any model system that permits genetic engineering, opening up spatially resolved analysis of embryonic development and other processes.

## Results

### Base editing can enable lineage recording with spatial readout

To ultimately capture detailed lineage relationships between cells while maintaining spatial context, we first sought to design an image-readable stochastic base editing system. One possible system would use designed target sequences that would be editable at a single base. For example, the A-to-G base editor ABE could target a set of defined sequences to stochastically edit each target site. However, this scheme is susceptible to convergent edits in unrelated cells, i.e. homoplasy. In the limit of complete editing, every editable A would be converted to a G in all cells, and no lineage information could be recovered.

To circumvent this issue, we used a modified design which takes advantage of the ability of the ABE to stochastically edit target sequences into one of multiple stable outcomes^24^. In our case, AA dinucleotide sequences in the target window are converted to any of three edited end-point states (GA, AG, or GG) (**Figure 1b**). Critically, because the GA and AG states each disrupt binding to the base editor gRNA, they are not expected to undergo further editing to GG. This dinucleotide editing scheme in principle reduces the likelihood of convergent editing, and prevents the effective “erasure” of recorded information at long times.

Based on this principle, we designed a library of barcoded, editable target arrays that could be integrated into the genome (**Figure 1c**). Each target array contained 6 tandem editable target sites, with unique protospacer sequences outside of the AA dinucleotide, so that each of the 6 target sites required a distinct gRNA sequence for editing (**Figure 1c**). The arrays were flanked with piggyBac inverted terminal repeats, to enable high-copy-number genomic integration. To distinguish different integrations from one another, we also incorporated two static (non-editable) random barcodes: first, a 10bp barcode (10^6^ variants), for compact readout by sequencing; second, a pair of static 80bp image-readable barcode sequences, each of which could take on one of 200 possible sequences, for a total diversity of 200^2^ = ∼10^4^ unique barcodes (see Methods for construction of plasmid libraries). Finally, to enable imaging-based “Zombie” readout of edits, the arrays incorporated a T7 promoter upstream of the editable array (**Figure 1c**)^23^.

To mutate targets at a tunable rate, and create the potential for signal recording, we built constructs that allow inducible expression of the ABE and gRNAs. ABE expression was placed under doxycycline (dox) control using the Tet-ON system, by stably integrating the reverse tetracycline-controlled transactivator (rtTA) (**Methods**)^25^. gRNAs were also made Wnt-inducible by expressing them from the 3’ UTR of an mTurquoise reporter gene under the control of a Wnt-responsive element^21,26^ (WRE). This promoter is active only in the presence of Wnt signaling ligands, and can be driven by the small molecule GSK-3 inhibitor CHIR99021 (CHIR)^21,26^. To generate fully functional gRNAs after expression, we flanked each gRNA with previously described ribozyme sequences^27^. After stable co-integration in mESCs using piggyBac transposition, we identified a clone, termed baseMEM-01, which contained 66 genomically-dispersed array copies with diverse static barcodes (**Figure 1d, Supplemental Figure 1**). This cell line, which allowed recording in live cells with imaging-based readout of barcode base edits, static barcode sequences, and endogenous gene expression using multiple rounds of hybridization and imaging^2–4^ (**Figure 1e**), was used for all subsequent experiments.

### Induction drives editing into diverse mutational states

Since lineage recording depends on edits being accumulated on the timescale of cell division, we next sought to analyze inducible base editing using the system. We exposed baseMEM-01 mESC cultures to the inducers CHIR, doxycycline, or both, over an 8-day period, a timescale long enough to allow multiple stem cell generations and, in an embryonic context, approach gastrulation. We collected samples at multiple time-points (**Figure 2a**). We then analyzed the editable barcodes by next generation sequencing.

**Figure 2:**
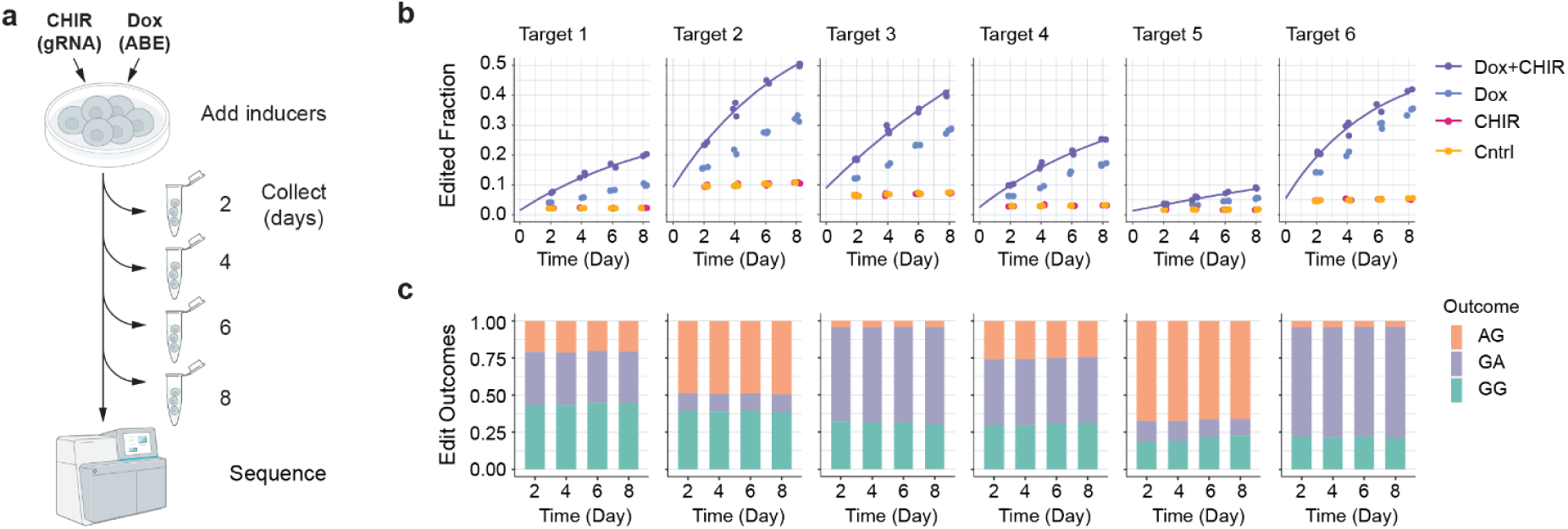
Dinucleotide targets accumulate edits over time in engineered mESCs. **(a)** Next generation amplicon sequencing quantified editing over time after induction of gRNAs and ABE. **(b)** All targets edited over time in the presence of the two inducers together, although at distinct rates **(b, purple)**. Dox induction alone drives editing at a slower rate **(b, blue)**. In the absence of dox, editing does not proceed at an appreciable rate **(b, red and gold)**. Three biological replicates were collected for each time point. The solid purple line shows the fit for a probabilistic model of editing over time (**Methods**). **(c)** Each target has a unique distribution of editing outcomes that remains constant as editing progresses.

Edits accumulated at all 6 target sites (**Figure 2b**). Without induction and in the absence of dox, some editing was observed. However, the fraction of such background edited sites generally remained constant during the time course, consistent with transient background editing during cell line construction, but minimal basal editing in stable clones. By contrast, in the presence of both CHIR and dox, edits accumulated rapidly at a rate that was well-fit by a model of editing with distinct edit rates at each site (**Figure 2b**, solid lines). Interestingly, in the presence of dox alone, editing still occurred, albeit at an attenuated rate. This non-zero edit rate could be due to basal transcriptional activity of the WRE promoter or to activation of gRNA expression by low levels of endogenous Wnt signaling in the mESCs. However, these results showed that doxycycline, with or without CHIR, could provide tight control of edit rate, making the line sufficient for lineage recording.

Next, we analyzed the distribution of edit outcomes at each site. Different sites exhibited distinct ratios of edit outcomes (**Figure 2c**). For example, site 6 exhibited a strong bias towards GA, and relatively little editing to AG. By contrast, sites 1 and 4 were more uniform in their outcomes. These differences in outcome bias across target sites are likely driven by differences in the sequences surrounding the dinucleotide for each target, which is known to impact CRISPR-Cas9 cleavage^28^ and base editing^24^. Regardless of how edits were biased, however, all target sites exhibited constant biases over time, consistent with the notion that bias is an intrinsic feature of the sequence context (**Figure 2c**). Most importantly, this consistency indicates that the GA and AG edit outcomes are stable for at least 8 days and do not become further edited to GG over the timescales of these experiments. These results thus validate the design goal of using inducible base editors to produce multiple distinct, individually stable states.

### Imaging recovers edited barcode sequences

We next turned from editing to imaging readout. In situ barcode readout creates the opportunity to assay lineage relationships without disrupting spatial relationships among cells. We developed an assay to readout barcode sequences through multiple rounds of single molecule RNA FISH (smFISH). We cultured BaseMEM-01 cells for 3 days — long enough for several cell generations — under editing conditions (3 µM CHIR, 1 µg/mL dox). We then fixed cells and performed in situ T7 transcription of barcodes using the previously described “Zombie” approach^23^ (**Figure 3a**, upper and lower left panels). Next, we hybridized pools of 24 primary probes designed to bind to one of the 4 possible dinucleotide states in each of the 6 target sites (**Figure 3a,b**). During hybridization, we also included 3 primary probes for each of the 400 different 80bp potential static barcode sequences, for a total of 1200 probes altogether (**Figures 1c, 3a,b**). We also included additional primary probe sets to analyze 12 different endogenous mRNAs, known to distinguish mESC pluripotency states in serum-LIF media^2,29^ (**Figures 3a,c**) (**Methods**).

**Figure 3:**
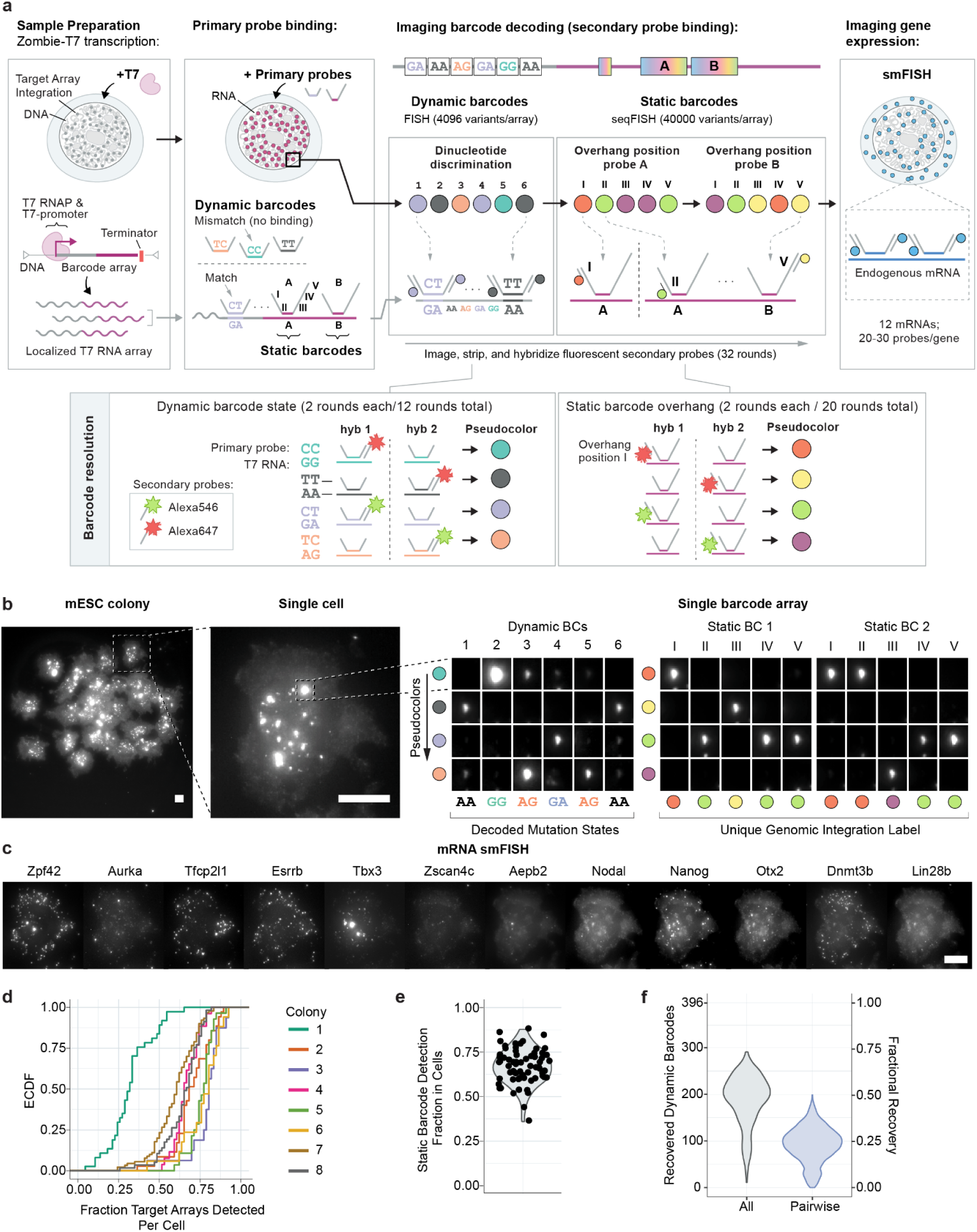
Multiple rounds of Zombie-FISH recover dynamic and static barcode states. **(a)** Barcode states can be recovered across multiple rounds of microscopic imaging. Ectopic application of T7 polymerase generates localized RNA clusters. Primary DNA probes are bound to the dynamic and static barcodes as well as to endogenous transcripts, competing primary probes against each other for binding to the different possible dynamic barcode variants. Each primary probe has an overhang sequence allowing for binding of one or more fluorescently labeled secondary probes, which are hybridized, imaged, and stripped away sequentially to recover barcoding **(b)** and transcriptional **(c)** information. **(d)** Across 8 colonies, we recovered 50-80% of target arrays per cell. One colony had dramatically lower barcode recovery and was excluded from further analysis (**d**, colony 1). **(e)** Each unique target array is recovered in a similar fraction of cells. **(f)** We recovered approximately 200 dinucleotide dynamic targets with high confidence per cell, with around 100 of these measured jointly between any pair of cells. Scale bars are 20 µm.

After hybridizing all primary probes, we sequentially added sets of fluorescently-labeled secondary probes, designed to hybridize to corresponding “overhang” sequences engineered in the primary probes (**Figure 3a**, lower panels). For each set of secondary probes, we imaged cells in all channels, and then stripped secondary probes to enable the next round of secondary hybridization and imaging. We used two orthogonal fluorescent channels to halve the number of rounds of hybridization required (**Methods**). We also labeled membranes with dye-conjugated wheat germ agglutinin to enable cell segmentation. All imaging was performed on a wide-field fluorescence microscope equipped with an automated fluid handling system similar to those described previously^2–4^. This procedure, similar to that used for seqFISH and related approaches^2–4,21–23,30–33^, allowed us to systematically probe dynamic barcodes, static barcodes, and endogenous genes over a total of 50 rounds of hybridization and imaging.

In the resulting image sets, individual target arrays could be identified as bright spots across multiple rounds of imaging. Most dynamic barcodes and static barcodes could be uniquely identified by a pseudocolor or set of pseudocolors (rows in image grids, **Figure 3b**). We developed a computational pipeline to detect target array spots and classify the barcode states within each target array (**Supplementary Figure 2**). Across 8 mESC colonies, we were able to detect roughly 50-80% of the 66 uniquely integrated target arrays in any given cell (**Figure 3d**). One colony had many fewer arrays detected than the others and was excluded from subsequent lineage analysis. The fractional detection of each unique barcode integration among all cells was broad but unimodally distributed, consistent with noisy detection efficiency in the absence of strong systematic differences among integration sites (**Figure 3e**). Overall, after applying quality controls, we detect roughly 200 high confidence dynamic characters, i.e. editable dinucleotides, per cell (**Figure 3f**). Of most direct relevance for lineage reconstruction, with these detection statistics, ∼100 shared characters could be confidently recovered in both members of any cell pair.

### Simulations show that baseMEMOIR can accurately reconstruct detailed lineage trees

We next asked how the depth of tree reconstruction depends on the distribution of shared characters between cell pairs, as well as other parameters, such as the rate and uniformity of cell divisions, the rate of editing, and the duration of recording. To address these questions, we simulated barcode recording and recovery. During the recording phase, we simulated editing within a single initial cell and its progeny for up to 12 cell generations (**Figure 4a, left**), setting barcode edit rates based on our time course editing dataset (**Figure 2b,c**). To simulate incomplete recovery of edit patterns, we stochastically subsampled the resulting barcode sequences corresponding to the observed empirical recovery distribution after hybridization and imaging (**Figure 3f**). As a result, different cell pairs shared different fractions of recovered barcodes (**Figure 4a, middle right**). Finally, we explored a filtering strategy to improve reconstruction accuracy (at the cost of reduced numbers of cells per tree) by restricting analysis to cell pairs with a minimum number of shared recovered barcodes (**Figure 4a, right**).

**Figure 4:**
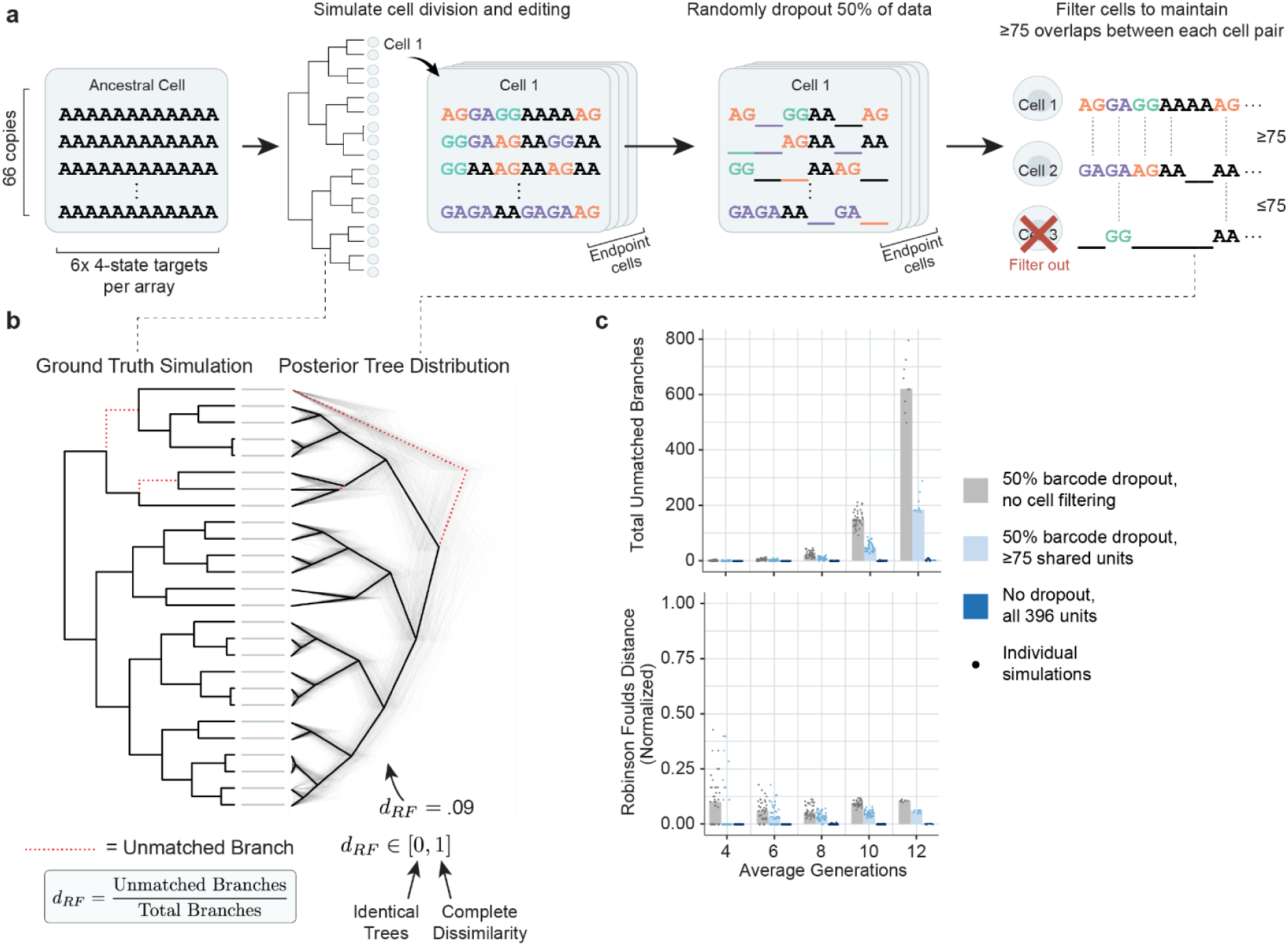
Lineage can be accurately reconstructed for at least 12 generations in simulation. **(a)** To estimate the expected accuracy of reconstruction, we simulated cell division and stochastic editing starting with unedited barcodes, represented as sets of AA dinucleotides (**left**) over time to produce heterogeneous edit patterns. We then either retained all sequences or dropped 50% of the data to represent random FISH detection losses, and filtered out cells that had few barcode characters overlapping with those measured in other cells (**right**). **(b)** Based on these ground truth simulations, we reconstructed lineage relationships and computed the Robinson-Foulds distance between the ground truth input (**left**) and reconstructed output (**right**) trees. **(c)** Reconstruction accuracy was nearly perfect without barcode dropout (**dark blue dots**). With dropout, we observed ∼ 10% error rates with tree depths up to 12 cell generations (**c, gray dots**). In the presence of dropout, filtering cells with few shared units moderately improved the reconstructed tree (**c, light blue dots**).

Next, we reconstructed lineage trees and compared them to the ground truth trees from the forward simulations (**Figure 4b**). As a metric of reconstruction accuracy, we used the normalized Robinson-Foulds distance, which quantifies the fraction of unmatched branches between the ground truth and reconstructed trees. For reconstruction, we adapted the Bayesian BEAST2 phylogenetic reconstruction framework, by incorporating a custom base editing model^34^ (**Methods**). BEAST2 uses Markov Chain Monte Carlo (MCMC) sampling to estimate the posterior probability distribution over different tree topologies and other system parameters. Briefly, it samples a forest of possible trees in proportion to their probability density (**Figure 4b**). As a Bayesian method, it allows for model-based inference, explicitly incorporates prior knowledge, and quantifies uncertainty in reconstruction.

In the ideal case of full recovery of all barcode edits in all cells, we obtained near perfect recovery of full lineage relationships for trees up to 12 cell divisions deep (**Figure 4c**). When ∼50% of barcodes were lost, error rates for the same 12 generation tree increased to ∼10%. However, this error rate could be reduced by restricting analysis to cells sharing at least 75 jointly measured barcode positions (**Figure 4c**). In contrast to the simulations, the experimental system could introduce additional factors such as errors in barcode readout, variability in mean edit rates between cells, and pre-existing edits in the ancestral (root) cell. Nevertheless, these simulations suggest that baseMEMOIR, with empirically observed error and edit rates, should be capable of reconstructing multi-generation lineage trees with cell cycle resolution at ∼90% accuracy.

### BaseMEMOIR reconstructs lineage trees in mESC colonies

mESCs are known to undergo spontaneous reversible transitions among a set of molecularly and functionally distinct cell states, ranging from 2C-like to formative and primed epiblast-like states, in serum-LIF conditions^2,29,35–40^. A fundamental question about the mESC state-switching process is the structure of the transition graph, i.e. which transitions occur and at what rates, and how those rates are influenced by input signals. In particular, CHIR, a Wnt pathway agonist that is often used to maintain pluripotent cells, could impact the observed cell states and their transitions. Although CHIR is known to promote expression of key pluripotency genes and self-renewal in this system^36,37,41–47^, the effects of CHIR on single cell state transition dynamics remain unknown.

To study these dynamics, we grew mESC colonies over a 3-day period in the presence of CHIR and Dox, which also serve to induce editing (**Figure 5a**). After 3 days, we imaged the colonies as described above and recovered barcode states for 7 colonies out of 8 total measured (**Figure 3d**). In addition to reading out barcode states, we also recovered gene expression levels for 12 pluripotency state markers, then clustered to identify 5 gene expression states (**Figure 5b,c** **and Supplemental Figure 4**). We identified 3 major cell states comprising: 2C-like cells with high expression of Zscan4C; naive cells expressing transcription factors Nanog, Esrrb, and Zfp42; and formative cells that express Otx2 and Dnmt3b in the absence of naive pluripotency factors and Zscan4C (**Figure 5b**). Naive cells exhibited a distribution of gene expression levels that varied from more 2C-like to more formative (**Figure 5b,c** **and Supplemental Figure 4**).These states largely correspond to those described in previous work^2,29,40^, although we observed high and relatively homogeneous Tbx3 expression across all cell states, in contrast to observations of cells grown in serum/LIF without CHIR^2,29^. This difference is consistent with previous work showing that Tbx3 is significantly regulated by CHIR in the observed direction^47^.

**Figure 5:**
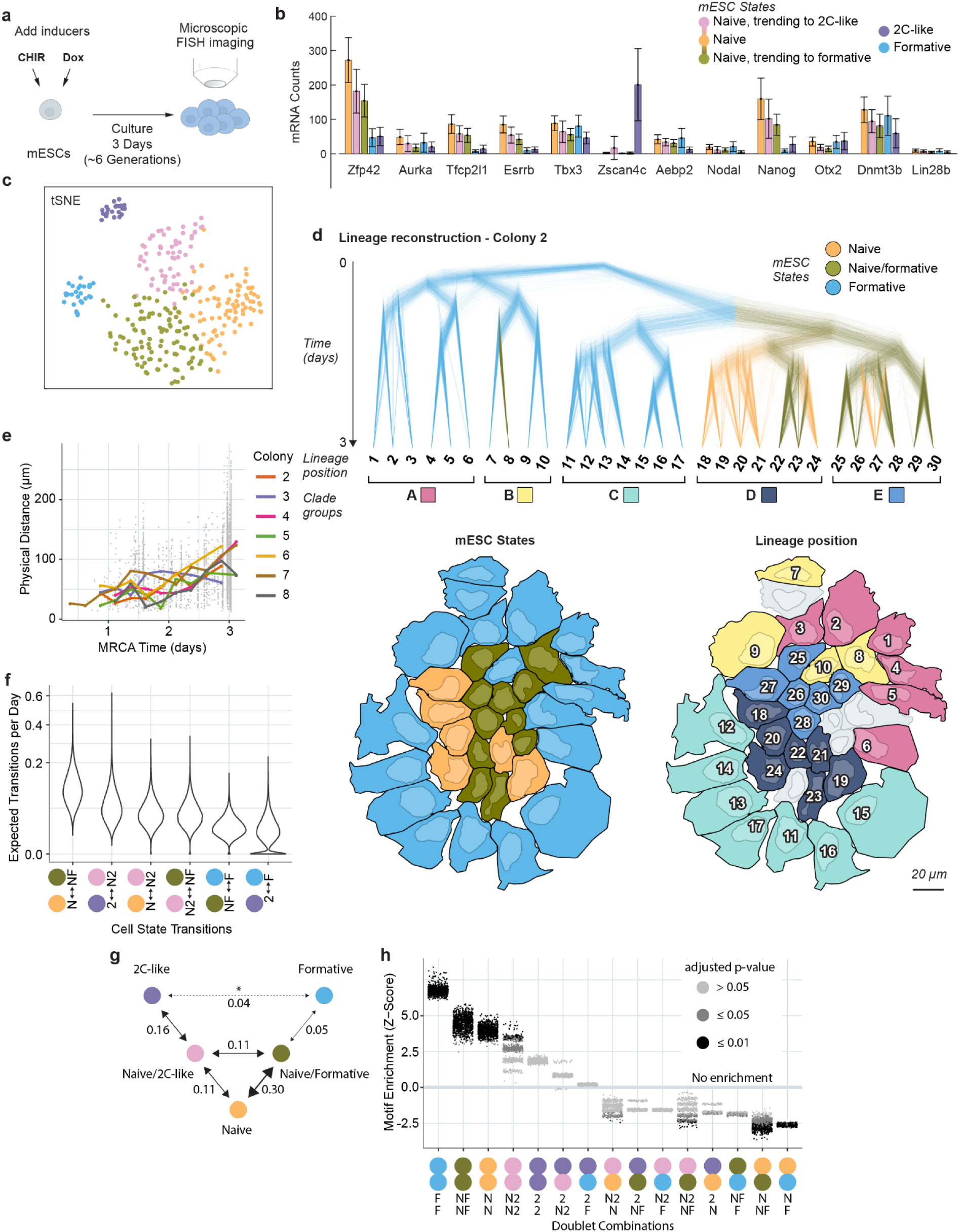
Joint measurements of lineage, gene expression, and spatial position reveal cell state transition dynamics. **(a)** We recorded lineage relationships in mESC cells cultured in serum-LIF media over a 3 day period, inducing editing with 3 µM CHIR and 1 µM Dox. **(b,c)** Cells clustered into 5 states based on gene expression as measured by smFISH. Two clusters were well separated from the other groups while three clusters appeared continuously related and expressed different levels of key marker genes (**see Supplementary Figure 4**). **(d-g)** Lineage reconstruction infers topological lineage tree relationships, cell division timing, ancestral cell states, and transition rates between those states. Uncertainty in lineage tree measurements is visualized by overlaying trees sampled from the posterior distribution of trees generated by Markov chain Monte Carlo for each colony (**d, top; Supplemental Figure 5**). Cell states and clade groups from the lineage tree can be mapped to the spatial colony images to qualitatively inspect the relationships between cell state, lineage, and spatial location (**d, bottom; Supplemental Figure 5**). **(e)** Spatial distance is larger between cells with more distant common ancestors. **(f)** Several cell state transitions were inferred to have nonzero median values across all posterior samples. **(g)** These state transitions predict a restricted cell state transition graph. One transition (denoted by *) contained a high fraction of posterior samples with a transition rate of 0. Numbers indicate the median expected number of transitions per day for cells of the given type. **(h)** Several doublet motifs are significantly over or underrepresented across the lineage tree posterior samples. N: Naive; 2: 2C-like; F: Formative; N2: Naive, trending to 2C-like; NF: Naive, trending to formative; MRCA: Most recent common ancestor.

We next applied the BEAST2 system described above to reconstruct lineage trees for cells in these colonies. To incorporate information of cell state and spatial position, we extended the underlying editing model (**Figure 4**) to represent these additional cellular properties, and thereby allowed simultaneous inference of cell state transitions and spatial movement alongside cell lineage (**Methods**). Applied to the mESC colonies, this approach reconstructed lineage relationships and division times with relatively low uncertainty in most cases, as indicated by the limited “fuzziness” of the reconstructed trees (**Fig. 5d** **and Supplemental Figure 5**). However, there were some ambiguities in reconstruction. For example, in Figure 5D, cell 23 is roughly equally likely to be a sister of cell 22 or cell 24. This illustrates the way in which uncertainties in the Bayesian reconstruction still provide specific alternative hypotheses rather than numerical confidence values. Additionally, in some cases, we did not capture all neighboring cells, which may introduce branch lengths longer than a single cell division (for example, see clades A and B of colony 7, **Supplemental Figure 5**). Together, these results demonstrate that the recording system allows precise lineage reconstruction of 3-day clonal mouse ESC colonies, with tree sizes of 30-50 cells.

The reconstructions also allowed estimation of cell state transition dynamics. To constrain the transition model, we treated transitions as a reversible, symmetric, continuous time Markov process (see **Methods** for discussion of these assumptions). Of the 10 possible symmetric interactions, posterior estimates suggested that ∼5 occurred at appreciable rates during the growth of these colonies (**Figure 5f,g**). A sixth transition, between 2C-like and formative states, was suggested by a single event in colony 7 (**Supplemental Figure 5**). This event is unexpected given previous inference of chainlike dynamics, with these two states at opposite ends^29^, however it could be a result of CHIR exposure, which was not present in previous work. Further, a substantial amount of posterior probability indicates a negligible rate for this transition; more data would be needed to clarify this result (**Figure 5f, Supplemental Figure 5**). Overall, these reconstructions suggest frequent (median of 0.15 transitions per day across all colonies) transitions among Naive/2C-like, Naive, and Naive/Formative states, and frequent conversion between Naive/2C-like and 2C-like states (**Supplemental Figure 5**). Interestingly, the inferred transitions correlate with expectations based on transcriptional similarity among states, even though this information was not provided to the model (**Figure 5b,c** **and Supplemental Figure 4**).

### BaseMEMOIR recovers lineage relationships, cell states, and spatial relationships in mESC colonies

By mapping lineage trees back onto the original images, we were able to simultaneously observe spatial and lineage organization of colonies (**Figure 5d**, lower panels). This analysis revealed correlations between lineage history, spatial position, and cell fate. For example, in Figure 5d, the related D and E clades contain Naive and Naive/Formative cells, and were located towards the interior of the colony, while cells in clades A,B, and C were largely in the Formative state and located around the periphery. Individual cell morphologies also varied systematically, with cells in the periphery exhibiting larger sizes. Other colonies were less radially structured, but still showed strong correlations between spatial positions and lineage relationships within each colony (**Figure 5e, Supplemental Figure 5**). These results show that it is possible to impose lineage relationships on spatial colonies with cell state information.

Lineage motifs provide a complementary approach to analyzing cell state transitions^48^. They are defined as statistically over-represented patterns of cell fates on lineage trees, which reflect features of underlying stochastic cell fate control programs^48^. As a simple example, asymmetric division, in which sister cells acquire opposite fates, would be reflected in the enrichment of opposite fates among sibling pairs (“doublets”). In contrast to the inference of transition rates described above, lineage motifs can be identified with no assumptions about an underlying model. Applying Lineage Motif Analysis^48^ (LMA) to 1000 samples from the Bayesian posterior tree distribution, we identified 4 overrepresented doublet pairs with an adjusted p-value less than 0.05 for a majority of the posterior samples (**Figure 5h**). These cases involve siblings in the same state, consistent with infrequent state transitions. Siblings in the formative state were the most overrepresented, mirroring results from the Bayesian Markov model, which predicts the slowest transitions to and from the formative state (**Figure 5f,h**).

We also observe two statistically underrepresented heterogeneous sibling pairs (**Figure 5h**). The most underrepresented pair, containing naive and formative cells, was also qualitatively consistent with predictions of the Bayesian Markov model, which identified a negligible transition rate between these states. Additionally, the naive and naive/formative sibling pair was also significantly underrepresented. This corresponds to the most rapid inferred transition rate in the dataset (**Figure 5f**), consistent with high rates of independent transitions out of either the naive or formative states. Together, these results demonstrate how baseMEMOIR’s lineage reconstruction ability allows inference of lineage motifs.

Finally, we combined the lineage reconstruction with spatial and cell state dynamics to infer a property that would be difficult to analyze from sequencing-based readout or static snapshots alone: the relative spatial mobilities of different cell states. The inferred histories of cell state and spatial position can be visualized (**Supplemental Movies**). These movies represent one possible history based on a simple model of cell diffusion, taking the highest credibility inference from BEAST2. Together, these results show how spatial position, cell state, and lineage can be analyzed and reconstructed together, and used to infer features of cell histories.

## Discussion

A long-standing dream in biology is to image a tissue or organism and visualize not only its current state, but also its past history. Previous work has approached this ideal in different ways, including lineage recording by accumulation of irreversible recombination events and reconstruction of small trees, however these efforts were limited in the amount and scalability of memory storage^21–23^. Here, we introduce a new approach, baseMEMOIR, which provides much larger memory sizes and allows for deeper, more accurate lineage tree reconstruction, while preserving spatial structure.

To achieve this, baseMEMOIR introduces several key innovations. First, it uses base editors to introduce stochastic, but precise, edits at dense target arrays (**Figure 1c**). Second, it uses dinucleotide editable target sites, each of which can be edited to any of three permanent end states (**Figures 1b, 2c**). Third, to discriminate between those states we expanded the Zombie readout system^23^ to allow 4-way probe competition (**Figure 3a,b**). Fourth, baseMEMOIR massively expands the amount of memory accessible in single cells by incorporating 66 unique statically barcoded target arrays, collectively providing 792 bits of editable information in the baseMEM-01 cell line. Theoretically, this number could be readily increased with additional target array integrations without modifying other components of the system. Fifth, baseMEMOIR achieves high density recording, while maintaining compatibility with FISH-based readout of endogenous genes (**Figure 3c**). Finally, to address the challenge of lineage reconstruction from stochastic edits, we adapted the BEAST2 framework for Bayesian tree inference, both by adding a new mutation model and taking advantage of its phylogeographical and discrete trait models^34,49^ (**Methods**). We anticipate that this probabilistic framework should be applicable for a broad variety of lineage recording methods.

To demonstrate these capabilities, we applied baseMEMOIR to stem cells undergoing interconversion among transcriptional states^2,29,35–38,47^. This allowed us to reconstruct lineage trees for 7 colonies totaling 197 cells, with as many as 4-7 cell generations per colony (**Figures 5d, Supplemental Figure 5**). Further, we were able to infer transition rates for specific pairs of states. These rates were consistent with a role for Wnt (through CHIR) in influencing state dynamics relative to similar cultures in the absence of Wnt^2,29,47^ (**Figure 5f,g**). Future work could use baseMEMOIR to systematically compare the effects of different signals and perturbations on cell state dynamics. While probabilistic inference is not equivalent to direct time-lapse observation, it nevertheless is beginning to yield related insights that would ordinarily be concealed from any static endpoint measurements (**Supplemental Movies**). Extrapolating from the capabilities of this system to future implementations, such as those containing either more memory or linking signaling pathway activity to recording machinery^16,21^, it should become possible to infer increasingly detailed views of earlier dynamic events in complex multicellular settings, effectively “decorating” lineage trees with events, such as changes in cell state or even movements in space. BaseMEMOIR should also allow one to infer state-switching dynamics and developmental programs using approaches such as Kin Correlation Analysis and Lineage Motif Analysis that exploit lineage tree information^29,48^.

While powerful, baseMEMOIR has some limitations. Because it does not directly probe the states of cells at earlier time points, it cannot directly detect earlier states that do not appear in the endpoint measurement. Analyzing systems at multiple timepoints could help to avoid missing transient states. Additionally, cells that die or migrate away prior to measurement will be omitted from the tree and could confound estimates of variation in cell cycle durations in different lineages. Similarly, failure to recover sufficient barcodes from an individual cell could make it difficult to classify. This issue may be addressed by further technical improvement to barcode imaging.

baseMEM-01 can immediately be used to explore stem cell differentiation and early mouse embryogenesis, among other phenomena. Looking ahead, baseMEMOIR should be readily adaptable to diverse developmental and physiological processes. The constructs and system can be transplanted to additional cell types using standard methods, and potentially combined with readout of additional “multi-omics” information such as chromatin accessibility^2^. One can therefore anticipate augmenting spatial cell atlases with lineage information^1^, and using baseMEMOIR to investigate the role of lineage, signaling, and differentiation in disease progression.

## Methods

**Dynamic barcoding strategy.**

Dynamic barcode sequences consist of 20bp CRISPR target sites with 3bp downstream NGG PAM sequences. These were chosen by designing sequences with AA nucleotides at the location predicted to be edited by the ABE (positions 5-6 in the protospacer sequence)^50^, then screening them for significant, varied editing of the AA sites. 6 unique target sites are arrayed sequentially downstream of a T7 promoter sequence to enable imaging-based readout as described below (**Figure 1c**).

### Static barcoding strategy

Static barcodes consist of two variable 80bp sequences downstream of the 6 dynamic barcode targets (**Figure 1c**). A pooled plasmid library was formed by generating constructs with 200 variants at each of the two 80bp regions, for a total of 40,000 unique sequences (**Supplemental Data**). Each sequence contains three unique primary probe binding sites for signal amplification during FISH readout (**Supplemental Data**).

### Plasmid construction

Plasmids were constructed in piggyBac backbones for later transposase mediated integration into the genome. The inducible ABE plasmid was made by integrating a tet-responsive promoter (TRE3G, Takara Bio) and ABE 7.10^50^ (Addgene #102919) into a piggyBac plasmid^51^ with neomycin resistance. The Tet-On 3G protein gene used to activate the ABE in a doxycycline dependent fashion was supplied as a piggyBac plasmid with a pEF promoter and puromycin resistance.

The dynamic and static barcode arrays were constructed in a piggyBac vector containing hygromycin resistance and double T7-T3 promoter sites followed by the dynamic barcode array, which was synthesized by Integrated DNA Technologies (IDT). The static barcode was then integrated 3’ of the dynamic barcode array. The static barcode was composed of two sites of 80 bp each, with 200 possible sequences for each of the two sites to give an overall possible barcode diversity of up to 40,000 unique sequences. The static barcode sequences were synthesized by Twist Bioscience and amplified with the appropriate cloning ends by PCR. The 5’ primer for the first static barcode site had a set of 10 random nucleotides to provide a further NGS-readable ID to each barcode. A mix of Gibson and sticky end cloning were used for plasmid construction.

The plasmid library containing static barcodes was generated by transforming high-efficiency competent cells (NEB C3019), then plating them onto a large surface area of LB-agar (∼30 10-cm petri dishes) to generate a large number of colonies. These were scraped and pooled into a single liquid culture. Subsequently, plasmid DNA was collected using multiple DNA Miniprep columns (Qiagen 27104) and pooled.

An array of six gRNAs targeting the six sites of the dynamic barcode were integrated in the 3’ UTR of an NLS-mTurquoise gene. Each gRNA sequence was flanked by the hammerhead and HDV ribozyme sequences on upstream and downstream sides, respectively, in order to excise the gRNA from the transcript. These gRNA-ribozyme sequences were each synthesized as gBlocks by IDT and combined by assembly of unique sticky end junctions into the piggyBac plasmid. A Wnt-responsive promoter was integrated to drive expression of the mTurquoise-gRNAs construct. This plasmid included blasticidin resistance for subsequent mammalian selection.

#### Primary probe library construction

Primary probes for dynamic barcode readout were purchased from IDT as individual sequences. The primary probe library, containing 1200 probes targeting all static barcode variants across both regions (3 probes per variant, 200 variants per region, 2 regions), was ordered as an oligoarray pool from Twist Bioscience. Each probe was assembled with a 35-nucleotide sequence complementary to the static barcode sequence, five 15-nucleotide readout sequences uniquely labeling each variant separated by a 2-nucleotide spacer, and two flanking primer sequences to allow for PCR amplification of the probe library (structure 5’-(primer 1)-(readout 1)-(readout 2)-(probe)-(readout 3)-(readout 4)-(readout 5)-(primer 2)-3’). The probe library was amplified following an established protocol^2^.

Endogenous marker genes were selected based on previous work.^2,29^ Probes for non-barcoded sequential smFISH of gene markers were a kind gift from Long Cai, generated as described previously^2^, using a single readout sequence repeated four times in place of a unique barcode (structure 5’-(primer 1)-(readout 1)-(readout 1)-(probe)-(readout 1)-(readout 1)-(primer 2)-3’).

#### Readout probe synthesis

Fluorescently conjugated secondary readout probes 15-nt in length were designed as in previous work^2,33^. Probe sequences were ordered conjugated to AlexaFluor 546 or 647 from IDT as indicated (**Supplemental Data**).

#### Coverslip functionalization

24 x 60 mm coverslips were functionalized prior to cell culture. Coverslips were first rinsed in 100% ethanol, then dried and functionalized using a plasma cleaner on the high setting for 5 minutes. Coverslips were subsequently immersed in 1% bind-silane (GE, 17-1330-13) solution (1% bind silane, 10 mM acetic acid in 90% ethanol) for 1 hr at room temperature. Coverslips were rinsed in 100% ethanol then heat dried in an oven at 90 C for 30 minutes before being treated with 100 ug/mL Poly-D-Lysine in water overnight. The following day, slides were rinsed with nuclease free water and air dried. Slides were stored for up to 2 weeks at 4 C prior to use.

Just before cell attachment, coverslips were treated with UV in a biosafety cabinet for 5 minutes, then the surface was treated with 10 ug/mL laminin (Biolaminin 511 LN, Biolamina) at 37 C for 90 minutes. Laminin was removed, then cell suspension was added directly to the surface for attachment.

#### Cell culture

E14 mES cells (ATCC cat. No. CRL-1821) were cultured in medium containing GMEM (Sigma), 15% ES cell qualified FBS (Gibco), 1x MEM non-essential amino acids (Thermo Fisher Scientific), 1 mM sodium pyruvate (Thermo Fisher Scientific), 100 µM B-mercaptoethanol (Thermo Fisher Scientific), 1x penicillin-streptomycin-L-glutamine (Thermo Fisher Scientific) and 1000 U/mL leukemia inhibitory factor (Millipore). For cell engineering and standard culture, cells were maintained on polystyrene (Falcon) plates coated with 0.1% gelatin (Sigma) at 37 C and 5% CO2.

#### Cell line engineering

Sequences of all integrated constructs are reported as Supplementary Data. BaseMEMOIR components were integrated over several rounds of transfection and selection. For all transfection steps, mESCs were cultured in 24 well plates, then cotransfected with the plasmid(s) to be integrated as well as piggyBac transposase plasmid with HD FuGENE transfection reagent. First, cells were cotransfected with ABE and Tet3G activator plasmids. The cells were allowed to recover for a day, passaged, and then underwent selection with 400 ug/mL neomycin followed by 500 ug/mL geneticin. Cells were plated sparsely in a 10 cm dish to grow monoclonal colonies, and then the monoclones were selected and grown in 96 well plates. Clones were screened for ABE expression after dox induction by qPCR, then subsequently by FISH to identify clones with homogenous expression among single cells.

Barcode target plasmids were integrated into the parental line containing inducible ABE by a second round of transfection, then selected with 100 ng/mL hygromycin as previously described. Monoclonal colonies were selected as previously, then screened by qPCR for high relative copy number. Zombie-FISH (described below) was used to screen promising candidates and select the clone with the highest visible integration number.

Finally, gRNAs and additional ABE plasmid were integrated into the most promising line from the previous step. Cells were selected with 15 ng/mL blasticidin, then monoclonal lines were generated as described above. Clones with a clear mTurquoise expression upon addition of 3 µM CHIR, which indicated expression of the gRNA construct, were kept for further analysis.

The best clones were tested for array targeting by adding 1 ug/mL doxycycline and 3 µM CHIR for multiple days followed by Sanger sequencing. Editing resulted in mixed peaks at the edited bases. One of the clones (baseMEM-01) was identified to have the most editing via this approach and was used for all subsequent experiments.

#### Next generation sequencing

Genomic DNA was extracted from cells using the DNeasy Blood and Tissue Kit (Qiagen) according to manufacturer instructions. Amplicon libraries containing the dynamic barcode sequences and short NGS static barcodes were generated with a two-step PCR protocol to add Illumina adapters and Nextera i5 and i7 combinatorial indices. Indexed amplicons were pooled and sequenced on the Illumina MiSeq platform with a 600-cycle, v3 reagent kit (Illumina, MS-102-3003). Raw FASTQ files were aligned to a FASTA-format reference file containing the expected amplicon sequences. Alignment was performed using the Burrows-Wheeler alignment tool (bwa-mem^52^). Subsequent analysis and data visualization was performed in the R statistical computing platform, v 4.1.1^53^ (**Supplemental Data**).

#### Edit accumulation model

Edit accumulation at each target site was modeled by fitting Equation 1:

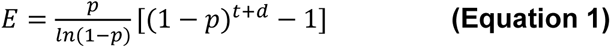

Here, edit accumulation, *E*, is a function of time, *t*, with parameters *p*, the probability of editing per unit time, and *d*, the duration of time during which edits accumulated prior to the zero time point, which accounts for empirically observed background edits (**Figure 2b**). This relation can be derived by assuming a probability *p* of a target site being editing per unit time *t* in a long string of target sites. After a unit of time *t*, we expect to see *p* edited targets and (1 − *p*) unedited targets. By the same logic, after another time step we expect *p* + *p*(1 − *p*) edited targets and (1 − *p*)^2^ unedited targets. After *T* time steps we would expect to see

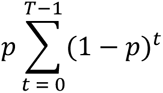

edited targets. Taking the limit of a discrete time step dt approaching zero, this sum can be approximated by the integral

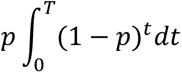

which simplifies to **Equation 1**.

Parameters were fit to editing time course data (**Figure 2**) to determine the empirical edit accumulation rate for each target using the “nls” function from the “stats” package^53^ in R (**Supplemental Data**).

**Stochastic simulations.** Barcode editing was simulated in R using the Gillespie method^54^ (**Supplemental Data**). Separate propensities were estimated for each editing outcome and target site by multiplying the edit accumulation parameter p (**Equation 1**) for each target site by the observed mean outcome proportion at each target site across time (**Figure 2c**). This stochastic simulation method recapitulates both the edit accumulation model fit and the empirical target state outcome distribution (**Supplemental Figure 3**).

Cell division was modeled by allowing editing until a predetermined cell division time, after which barcodes were duplicated before allowing editing to continue. Cell division waiting times were drawn from a distribution derived from Eyring-Stover survival theory that has been shown to model cell division times more accurately than the exponential distribution^55^.

Lineage relationships were reconstructed based on the resulting barcodes using BEAST2 software as described below, considering only barcode data (see BEAST2 XML files for complete modeling information, available at https://doi.org/10.22002/327t7-ke088).

Reconstructed trees were compared to simulated ground truth trees by computing the normalized Robinson-Foulds distance as implemented in the “RF.dist” function from the R package “phangorn”^56^.

### Zombie preparation

For Zombie and subsequent RNA-FISH, cells were plated on treated coverslips as described above. After culture and editing, coverslips were washed with 1 mL PBS with calcium and magnesium (PBS +/+) then fixed with a 1:1 solution of Methanol : Acetic Acid (MAA) for 20 minutes. RNase-free reagents were used for all subsequent steps to minimize RNA degradation. MAA was removed, then coverslips were transferred to a 100 mm petri dish and covered with 70% ethanol. Petri dishes were parafilmed and stored at -20 C to await imaging.

Immediately prior to imaging, coverslips were removed from cold storage and brought to room temperature. 70% ethanol was removed and replaced with a fresh solution of MAA, then incubated for 2 hrs at room temperature. The sample was washed twice with PBS +/+, incubating for 2-3 min between each wash. The final wash solution was removed and the sample was dried until all liquid had just evaporated. A custom fluidic cell, built to interface with a custom designed liquid handling system, was affixed to the coverslip surface^2–4^. Subsequent washes took place within the flow cell, manually adding reagents into the inlet of the cell and removing them from the outlet using a standard micropipette. The cells were washed with nuclease free water once, then replaced with T7 RNAP mix (New England Biolabs E2040S). The sample was incubated at 37 C overnight in a humidified tupperware.

The following morning, the T7 RNAP mix was removed and replaced with fresh T7 RNAP mix, then incubated for 1 hr at 37 C in the humidified tupperware. The mix was removed, then the sample was immediately fixed with 4% paraformaldehyde for 10 min. This solution was removed and the sample was washed three times with PBS +/+, then washed with 30% formamide probe wash buffer (30% formamide in 5x SSC with 9 mM citric acid, 0.1% Tween-20, and 50 μg/mL heparin, pH 6.0) for an additional 5 min. The wash buffer was replaced with primary probe hybridization mix, then incubated overnight at 37 C.

#### FISH imaging

Images were collected across multiple rounds of fluorescence hybridization to identify barcode and cell states. Formamide wash buffers and secondary probe hybridization mixes were generated immediately prior to imaging. A custom-built, automated liquid handling system was used to perform sequential rounds of in situ hybridization as previously described^2–4^. Briefly, the sample was connected to an automated fluidics system attached to a widefield fluorescence Nikon Eclipse Ti microscope. The custom-made automated fluid sampler was used to transfer readout probes in hybridization buffer from a 2.0 mL 96 well plate through a fluidic valve (IDEX Health & Science EZ1213-820-4) to the custom-made flow cell using a syringe pump (Hamilton Company 63133-01). Fluidics and imaging were integrated using a custom script controlling uManager. Eleven fields of view (FOVs), capturing 8 well separated regions of cell growth, were selected based on the DAPI signal. For each FOV, images were acquired with 0.5-micron z steps for twenty total slices. Integration of the automated fluidics system and imaging was controlled by a custom script written in uManager^57^.

First, twelve hybridization rounds were imaged to capture all dynamic barcode states. The hybridization buffer for each round included two unique 15-nucleotide readout probes (**Supplemental Data**) conjugated to either Alexa Fluor 647 (50 nM) or Alexa Fluor 546 (50 nM) in EC buffer (10% ethylene carbonate, 10% low molecular weight dextran sulfate, 4x SSC).

Probes were allowed to hybridize for 15 minutes. Excess probes were washed away with 10% wash buffer (10% formamide, 0.1% Triton X-100 in 2x SSC) incubating for 1 minute. Nuclei were re-stained with DAPI solution (5 ug/mL DAPI in 4xSSC) incubating for 2 minutes. The sample was washed with 4x SSC then imaged in antibleaching buffer (50 mM Tris-HCl pH 8.0, 300 mM NaCl, 2xSSC, 3 mM trolox, 0.8% D-glucose, 1000-fold diluted catalase, 0.5 mg/mL glucose oxidase). After imaging, readout probes were stripped off using 35% wash buffer (35% formamide, 0.1% Triton X-100 in 2x SSC). Although 55% formamide is typical for stripping readout probes, we used a lower amount to avoid stripping primary probes and losing signal, as our primary probes are shorter than normal for dynamic barcode rounds (only 20-nucleotides compared to 28). Images were collected after probe stripping to verify loss of signal. Due to occasional technical issues such as loss of focus during automated imaging, these twelve rounds were repeated a second time to collect backup images for each dynamic barcode round.

Static barcode sequences were captured by a similar scheme over twenty additional rounds of hybridization (see **Supplemental Data** for probe sequences), except using 55% formamide wash buffer to strip the readout probes. An additional six rounds of hybridization were used to capture the twelve gene markers described above. A final round of hybridization with wheat germ agglutinin (WGA) conjugated to Alexa Fluor 647 was used to stain cell membranes for downstream segmentation.

#### Image processing

Images were processed using custom Matlab scripts (**Supplemental Data**). DAPI signal was measured in each round of imaging and used to register images across each hybridization round. After registration, z-stacks were projected by their maximum intensity to yield one image per channel per hybridization round for each colony.

Transcribed barcodes form dots of variable intensity around the active site of T7 transcription. Dots were segmented using a combination of Laplacian of Gaussian filtering and watershed, requiring a maximum eccentricity of 0.8 to reduce noise. Loose parameters were chosen to detect all real dots at the expense of accepting some background noise. Binary images for each hybridization/channel were summed together to create a single mask, where pixel values represent the number of times a pixel was identified across all imaging rounds, termed “analog mask”.

Since each real dot should appear across all hybridization rounds in at least one channel, we further reduced noise by thresholding this image. We determined a threshold for each colony individually based on the elbow method. Frequently, we observed segmentation errors where the watershed algorithm was unable to separate adjacent dots from one another. We manually corrected these errors based on the analog mask, yielding a final binary segmentation mask of all detected barcode dots for each field of view.

We further generated DAPI segmentation masks using Ilastik to isolate individual cell nuclei^58^. Masks were manually corrected using ImageJ to separate nuclei that were segmented together. Cells that intersected the border of the image were excluded. Any Zombie dots identified outside of cell nuclei were filtered. Dots may not be completely captured by the binary mask within any given round of imaging. A K-nearest neighbors classifier was used to partition all pixels belonging to each cell to the nearest dot in the segmentation mask so that intensity values could be extracted.

Raw images were background subtracted to improve signal to noise. First, a tophat filter was used to globally reduce background. To further correct for variable intensity across images, local background correction was applied on a cell-by-cell basis by subtracting the median pixel intensity, excluding dilated segmented Zombie dots.

We extracted several features for each dot across all channels and hybridization rounds based on the background-subtracted raw images (total intensity; median intensity; 90^th^ percentile pixel intensity; pixel count; background median intensity; and intensity variance), taking the log + 1 of all intensity values.

We used a supervised machine learning approach to classify barcode states across each hybridization round. The barcode state is reflected in higher intensity fluorescence of probes that outcompete other possible binders (**Figure 3**). We manually classified approximately 1000 randomly sampled barcode spots for each image based on their pseudocolor intensities, then used this sample to train a support vector machine (SVM) classifier in Matlab (**Supplemental Data**). Some spots were ambiguous; these were omitted in manual classification. 10-fold cross validation was used to evaluate model generalization and control for overfitting (**Supplemental Figure 2**).

For dynamic barcode sites, we estimated the posterior probability for each spot belonging to each class under the SVM model. Many barcodes could be classified with high accuracy (>70% posterior probability, **Figure 3f** and **Supplemental Figure 2b**). For static barcode sites, class assignments were compared to the white list of possible barcode sequences. We filtered out Zombie spots with a character distance greater than 2 from an expected sequence and those which did not unambiguously correspond to a white listed static barcode, leaving 79.3% of all detected spots after filtering.

In many cases, duplicated barcodes were observed, where the same static barcode was identified multiple times in a single cell. These duplicates tended to be spatially localized and may be explained by either DNA replication or over-segmentation errors during analysis. For duplicated barcodes, classification probabilities were averaged at dynamic barcode sites and the most confident state was used for downstream lineage reconstruction.

Membrane masks were manually generated based on WGA staining images, then gene markers were identified using the bigFISH package dot detection method.^59^ Thresholds for dot detection were manually determined for each gene. Segmented spots, corresponding to mRNA molecules, were tallied within each cell as defined by the membrane segmentation mask. To validate the consistency of this method, we plotted the detection frequency for each gene across all cells that were measured in multiple images (**Supplemental Figure 6**). The measures were highly correlated.

### Cell type analysis

Cell types were determined by using k-means clustering on log transformed mRNA counts with 5 centers to group the most distinct sets of cells in the dataset. Dimensionality reduction by the tSNE method visualizes three groups as well separated and three of the identified cell states (Naive/2C-like, Naive, and Naive/Formative) as potentially a continuous distribution, although we note that dimensionality reduction techniques can obscure the true distances between cells and clusters in the higher dimensional transcriptome space. Most importantly, we identify unique allowed and forbidden transitions between each purported cell state through subsequent lineage analysis that is agnostic to the underlying transcriptional information, bolstering the claim that these five clusters of cells should be treated as distinct populations.

### Lineage reconstruction and Bayesian modeling

We used a Bayesian model under the BEAST2^34^ v2.7 framework that takes barcode information, end point cell labels, and cell centroid positions as input to jointly estimate lineage relationships, cell state transition dynamics, and cell motility. An XML file specifying all modeling information is provided as supplemental data and modeling choices are briefly described below.

Barcode information for each dinucleotide was extracted using Matlab and R scripts (**Supplemental Data**). Each of the four dinucleotide states (AA, AG, GA, and GG) was encoded in a single character (A, T, C, or G). Characters that were not recovered during imaging were marked as missing data by the “?” character. Cell division for each tree was modeled as a pure birth process (the Yule model) with birth rate estimated.

Barcode character mutation in our system is irreversible. With few exceptions^60^, existing BEAST2 packages only model reversible character transitions because these make computing tree likelihoods more computationally efficient. We developed a new irreversible character substitution model to better capture the evolutionary process that generated our data (available as the ‘irreversible’ package for BEAST2, with source code available from https://github.com/rbouckaert/irreversible). Under this model, each possible transition (AA to AG, GA, or GG) can take a unique rate value, which we assume is constant along the tree.

Stationary frequencies, which are used at the root of the tree to calculate the tree likelihood, are set at 1 for the AA state, and 0 for the others, reflecting our knowledge that every state is AA at the root of the tree. Since we know that targets can edit at different rates and into different outcomes, we allow the rate to vary across sites through the gamma site heterogeneity model, partitioning the allowable rates into 4 categories^61^. This model is shared across all trees. Furthermore, we use a strict molecular clock since we do not expect significant rate variation per branch.

Cell state transitions are modeled as a continuous time Markov chain with symmetrical transition rates possible between each state. These rates are assumed to be constant across time with an associated strict clock model. Transition rates are shared between all trees, so a single unified cell state transition model was estimated across all colonies. We assume symmetric transition rates based on previous work, which identified most transitions as roughly symmetric in this system^29^. In principle, the assumption can be relaxed, although it greatly increases the number of parameters in the model, thus increasing susceptibility to overfitting.

Cell motility was modeled as single parameter 2D diffusion along the surface of a sphere as implemented in previous phylogeographical work^49^. Spherical diffusion is a good approximation of diffusion in a 2D plane for small patches of the surface^49^ and its implementation is efficient. Accordingly, cell positions were mapped to geographical coordinates falling within 2 latitude and longitude degrees. The diffusion parameter describing motility was allowed to take unique values along each branch of the tree under a relaxed clock model.

Notably, all 7 colonies in this dataset were analyzed simultaneously under a single model. This allowed us to infer barcode character substitution and cell state transition models that are shared across all the data, reflecting our belief that all colonies are representative of the same underlying barcode mutation and cell state transition processes. We think this is reasonable given that all colonies are generated from a monoclonal culture grown in identical culture conditions over the same time period.

We chose priors to be uninformative with the exception of the root height, since we have strong prior information that experiments lasted 3 days. An uninformative but improper uniform distribution across all possible rates was chosen for barcode mutation rate, although this is not expected to affect the resulting analysis or MCMC mixing. Detailed prior information is recorded in the supplemental XML file for Figure 5 specifying all modeling choices.

Supplementary movies were generated by creating inferred still images of the maximum a posteriori histories of cells over time, incorporating inferred ancestral cell states, positions, and cell division timings (**Supplemental Data**). These still images were compiled into movies using the open-source video editor Shotcut (Meltytech).

### Lineage motif analysis

The posterior baseMEMOIR trees were analyzed using Lineage Motif Analysis (LMA) as described previously^48^, using the resample_trees_doublets, resample_trees_triplets, and resample_trees_quartets functions with 1000 resamples. These functions are available in the publicly available “linmo” package for Python (https://github.com/labowitz/linmo). To generate a z-score and adjusted p-value for all cell fate patterns across the entire posterior distribution of each tree dataset, 1000 synthetic datasets were generated by randomly drawing one tree from the posterior distribution of each tree dataset. Each synthetic dataset therefore contains 7 total trees. LMA was then performed on each synthetic dataset, and the distribution of z-scores and adjusted p-values was plotted for each cell fate pattern.

### Data availability

Raw image data is available at https://doi.org/10.22002/pmpby-gpj05. All analysis scripts, amplicon sequencing data, max projected image data, and additional supplementary files are available at https://doi.org/10.22002/327t7-ke088.

## Supporting information

All Supplemental Files

---

**Supplemental Figure 1:**
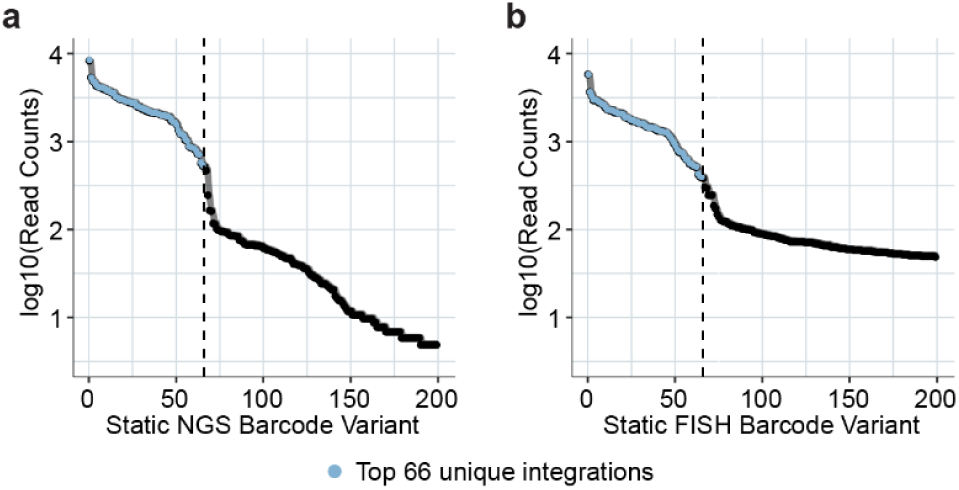
66 unique integrations are detected in the baseMEM-01 cell line. 66 barcode integrations were identified by next generation sequencing of target arrays amplified from genomic DNA. We quantified the number of reads corresponding to unique sequenceable **(a)** and image readable **(b)** static barcodes, identifying approximately 66 variants in each case. The top 200 most frequent variants are shown; we separated true variants from noise heuristically by identifying the “knee of the curve” (dashed vertical lines). Importantly, these 66 variants are all also identified in FISH experiments (Figure 3e).

**Supplemental Figure 2:**
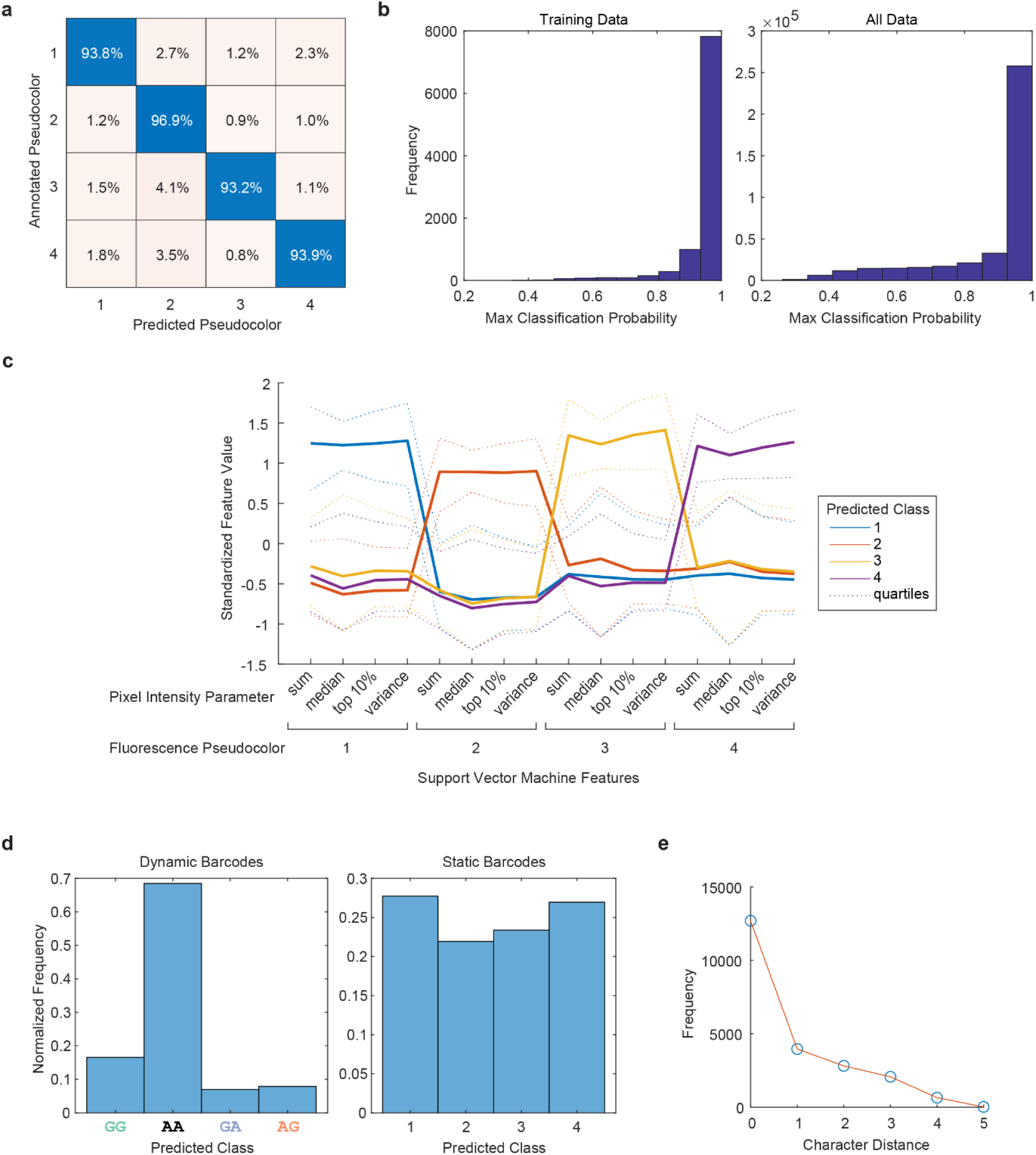
A support vector machine classifies barcode states based on fluorescence measurements. **(a)** Manually annotated barcodes are correctly classified by a quadratic kernel support vector machine (SVM) approximately 94% of the time. **(b)** Classification probability estimates are very high within the training dataset **(b, left)**. Outside of the training sample, most classification probabilities are still high but with a subset of predictions that are less certain **(b, right)**. The support vector machine predicts classes based on 16 fluorescence measurements corresponding to each pseudocolor as defined in Figure 3b. **(c)** Each class is well separated based on these features. **(d)** After 3 days of editing induction, many dynamic barcodes are identified as class 2, corresponding to the unedited state **(d, left)**. Static barcode classifications are more evenly distributed, as anticipated **(d, right)**. **(e)** Static barcodes decoded by FISH typically perfectly match the 66 image readable barcode sequences identified by sequencing (**Supplemental Figure 1b**), although a fraction of barcodes are recovered with one or more character differences relative to their closest match.

**Supplemental Figure 3:**
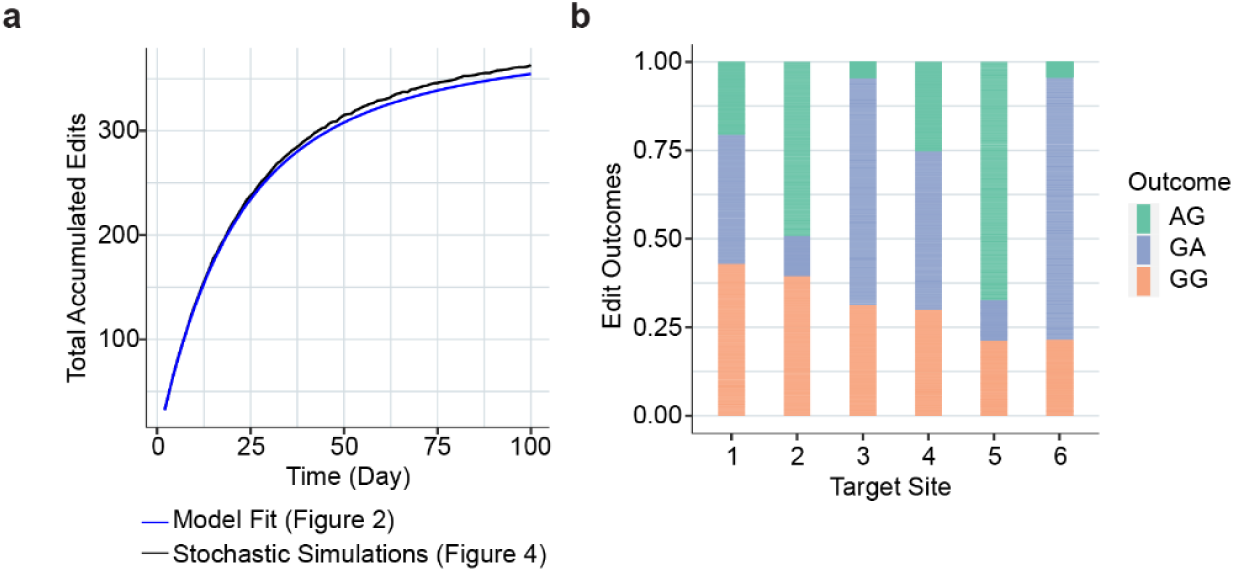
Stochastic simulations closely recapitulate the empirical editing process. **(a)** We developed a stochastic editing simulator based on the Gillespie algorithm that closely recapitulates the average edit accumulation model developed in Figure 2b (**Methods**). **(b)** The simulated edit outcome distributions for each target site match the observed distributions from Figure 2c.

**Supplemental Figure 4:**
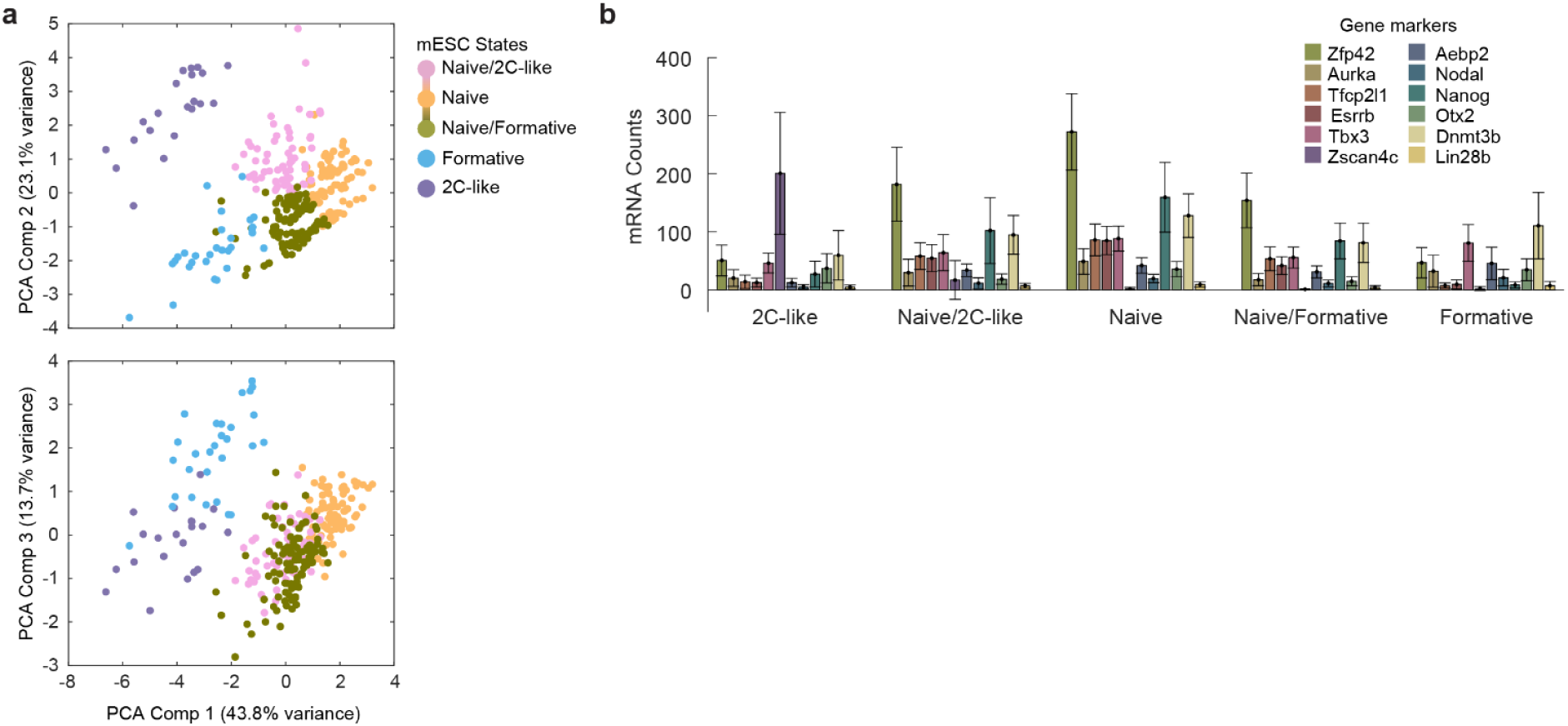
mESC gene expression clustering. **(a)** Principal component analysis is largely in agreement with nonlinear dimensionality reduction, with separation between major clusters observed along the first three components. The naive states also appear continuously related in this view. **(b)** Clusters have distinct marker gene expression patterns, with some similarity between the Naive/2C-like, Naive, and Naive/Formative states.

**Supplemental Figure 5:**
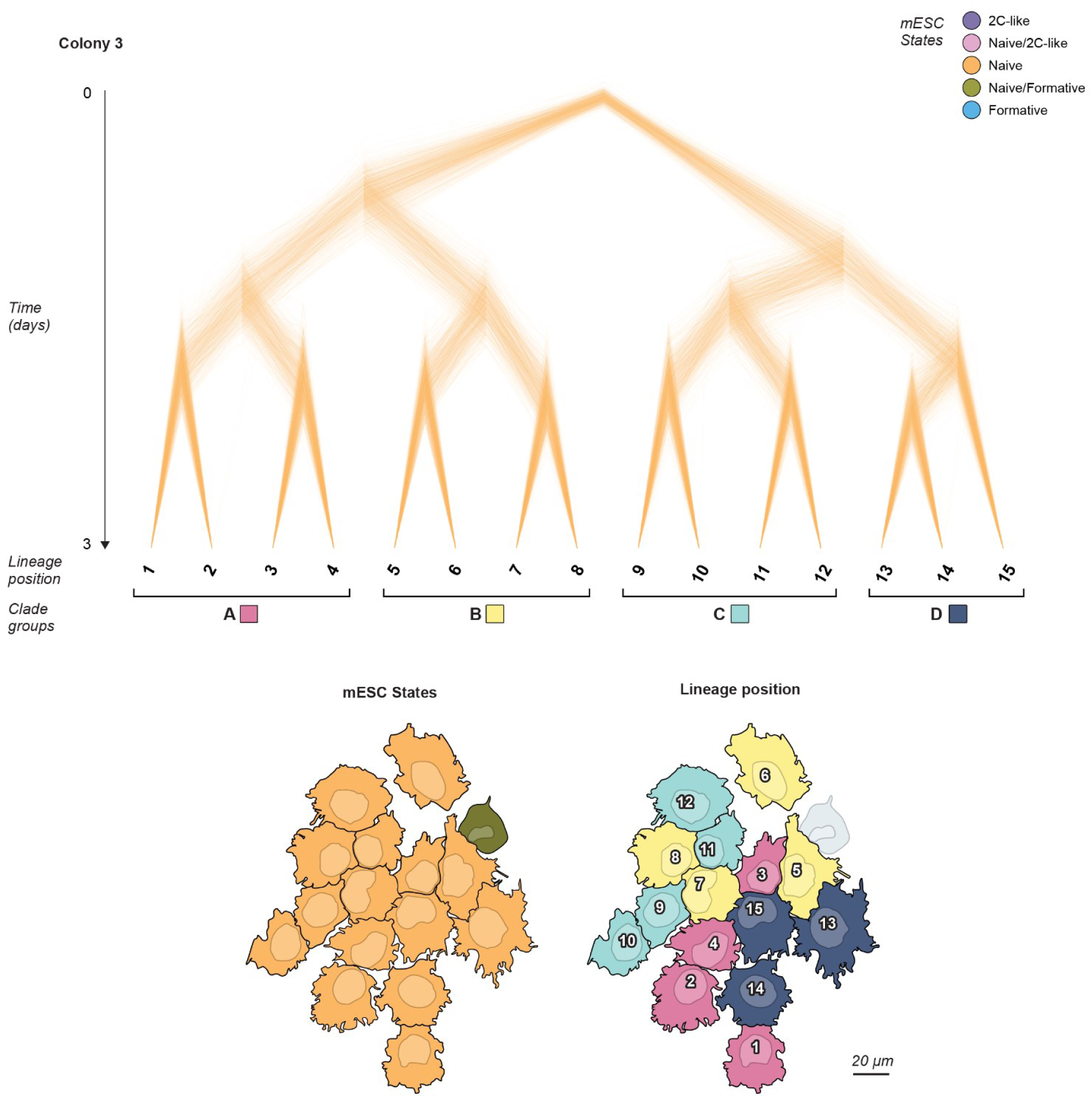
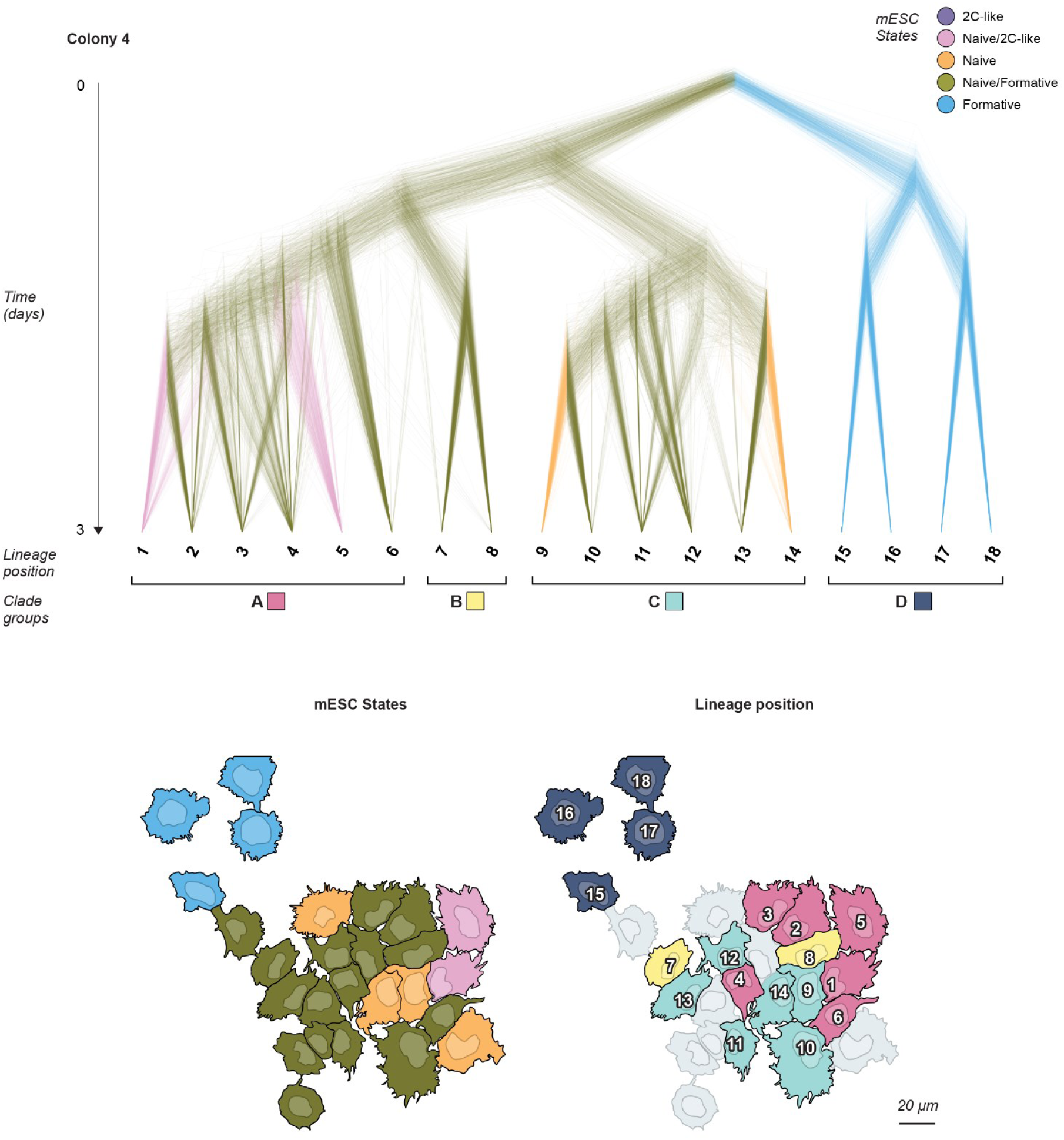
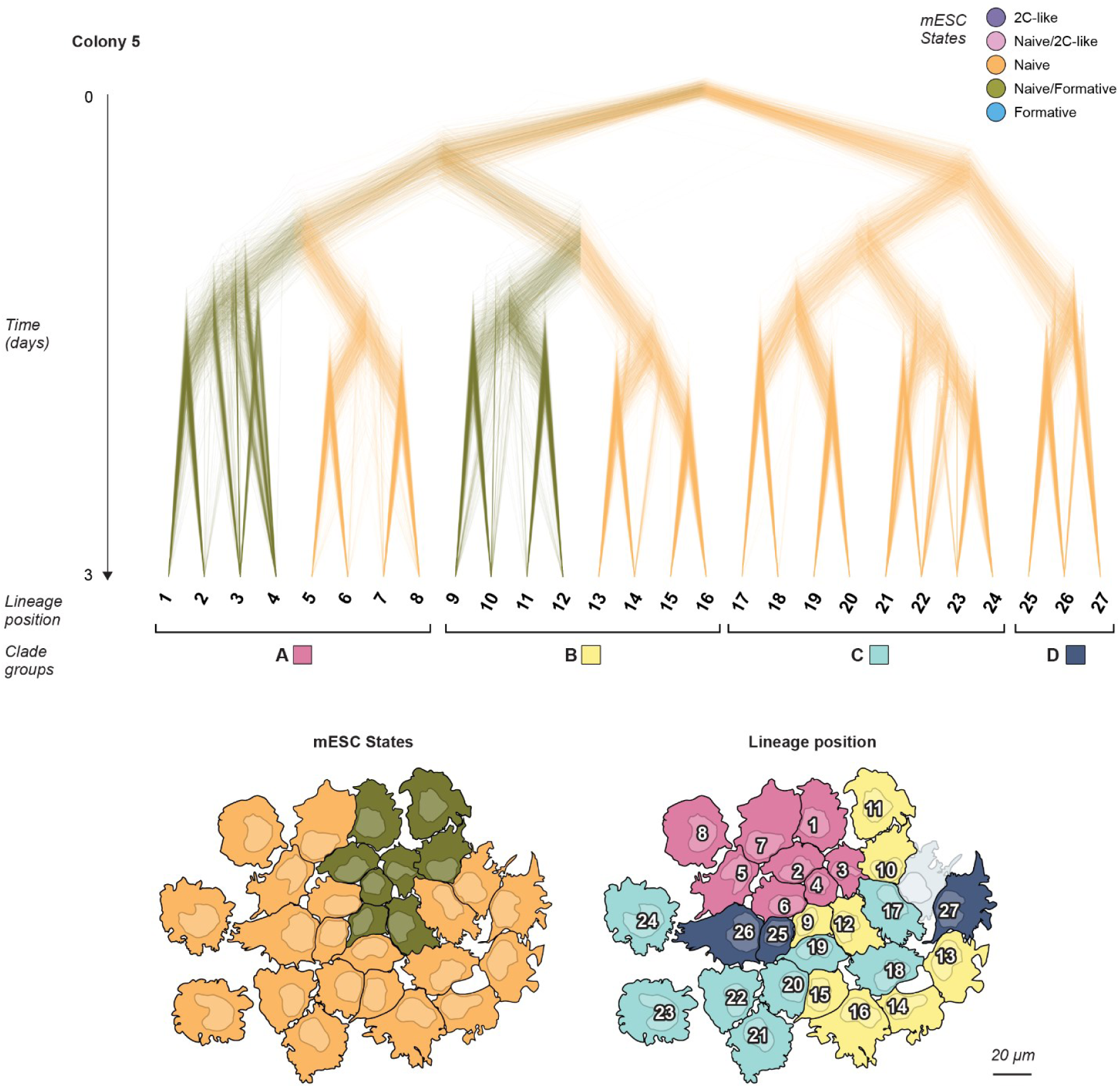
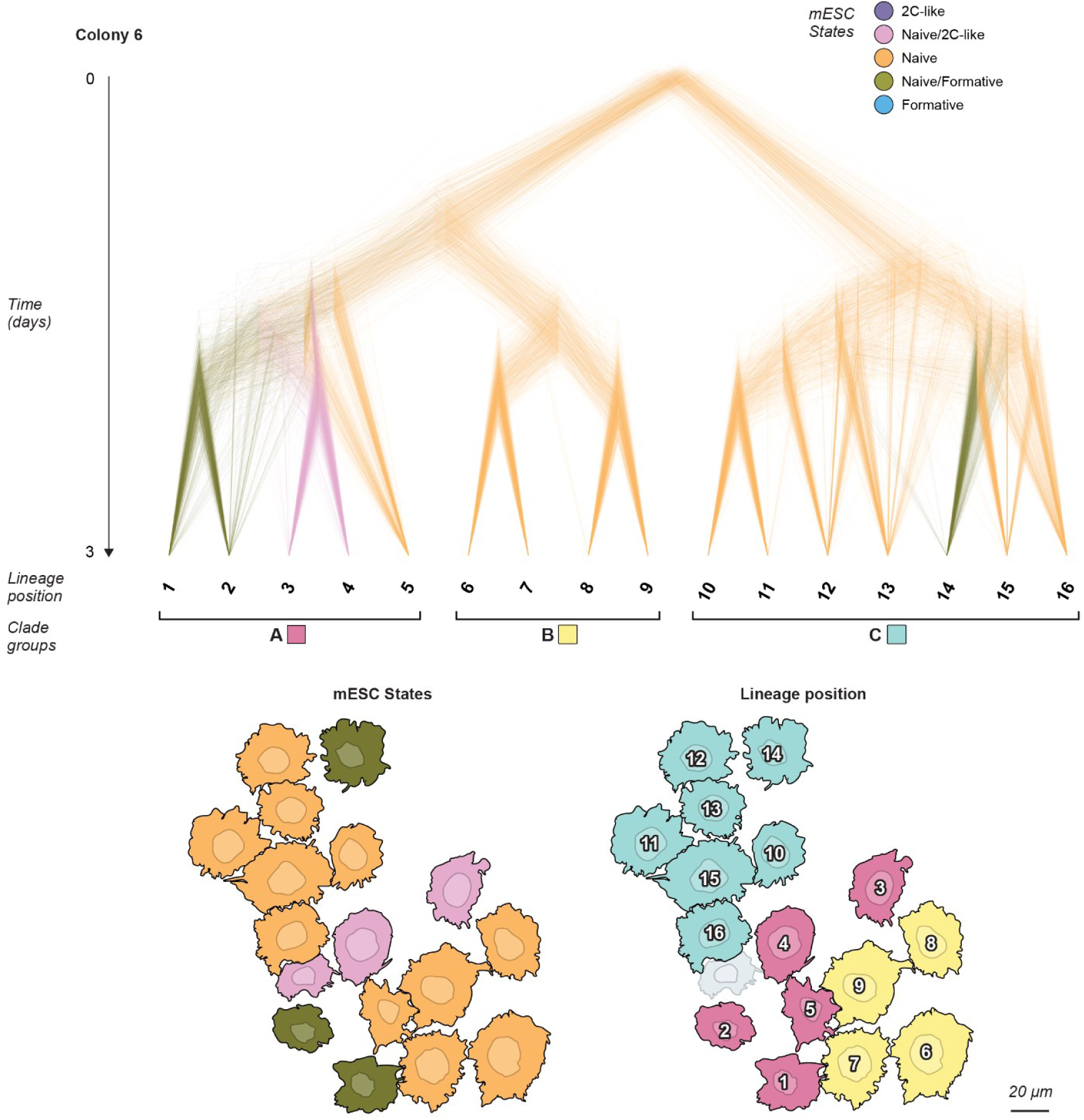
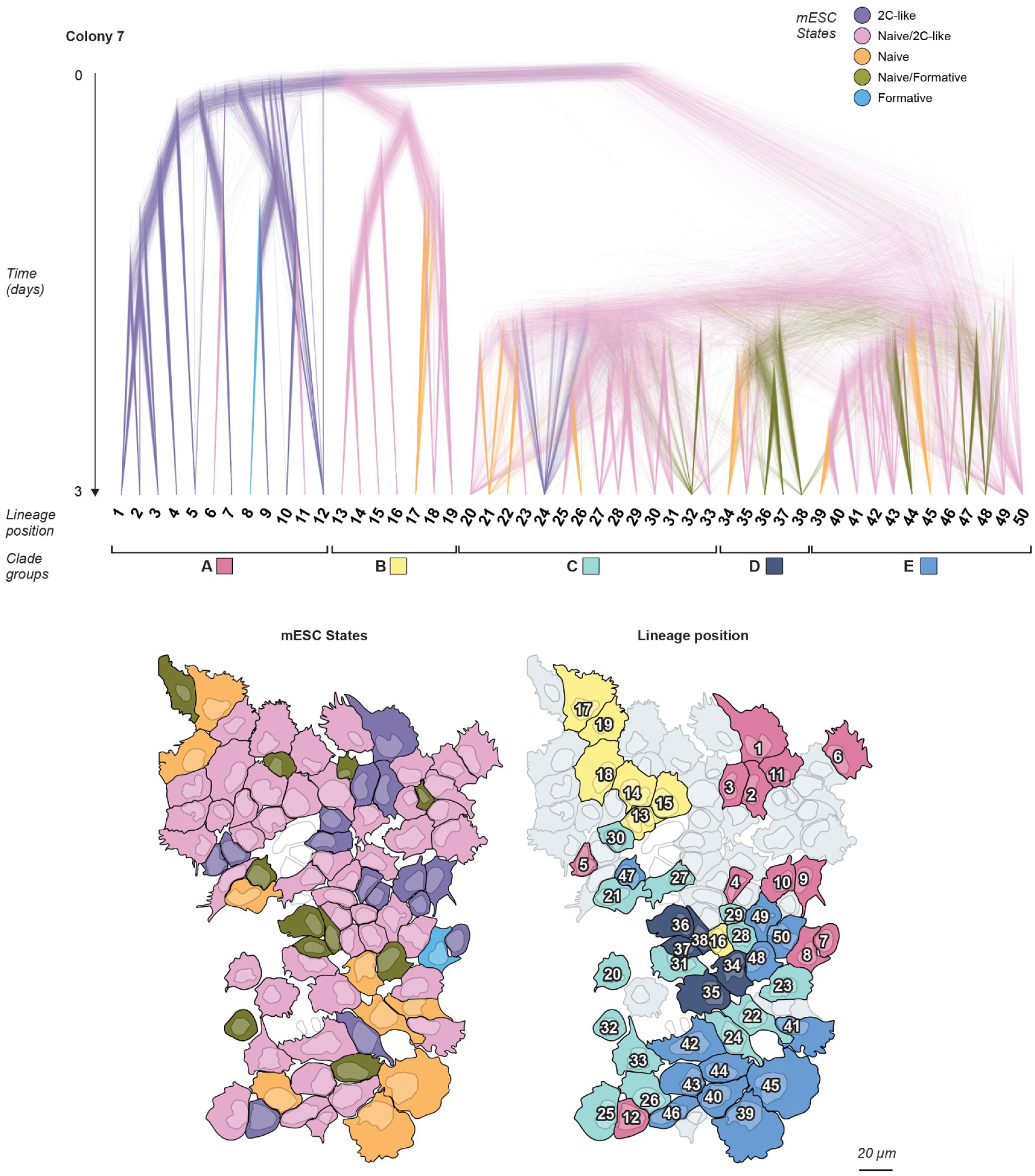
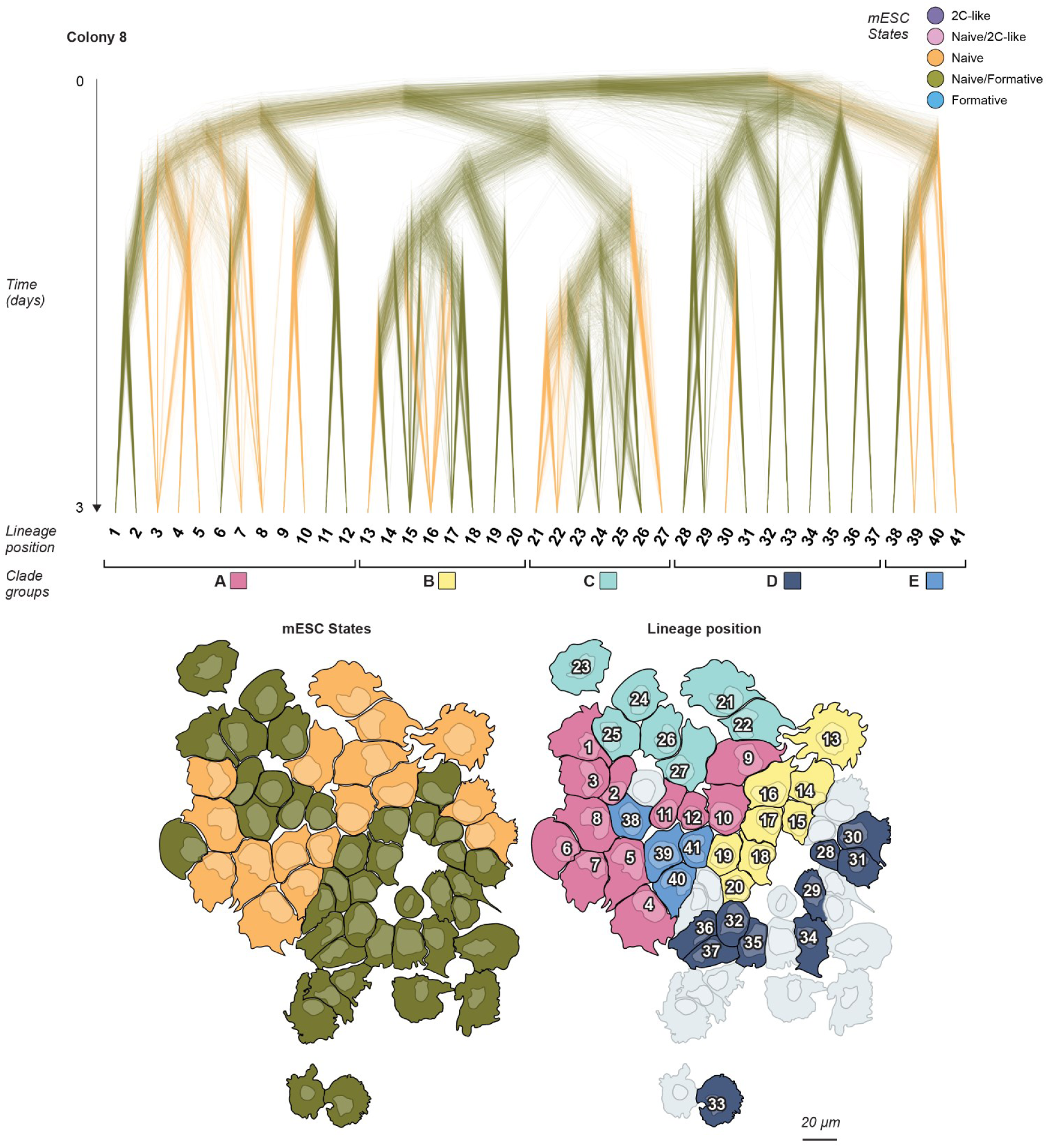
BaseMEMOIR reveals lineage relationships, cell states, and spatial positions across multiple colonies. Posterior tree distributions are visualized and mapped back to illustrations of each colony as in Figure 5d.

**Supplemental Figure 6:**
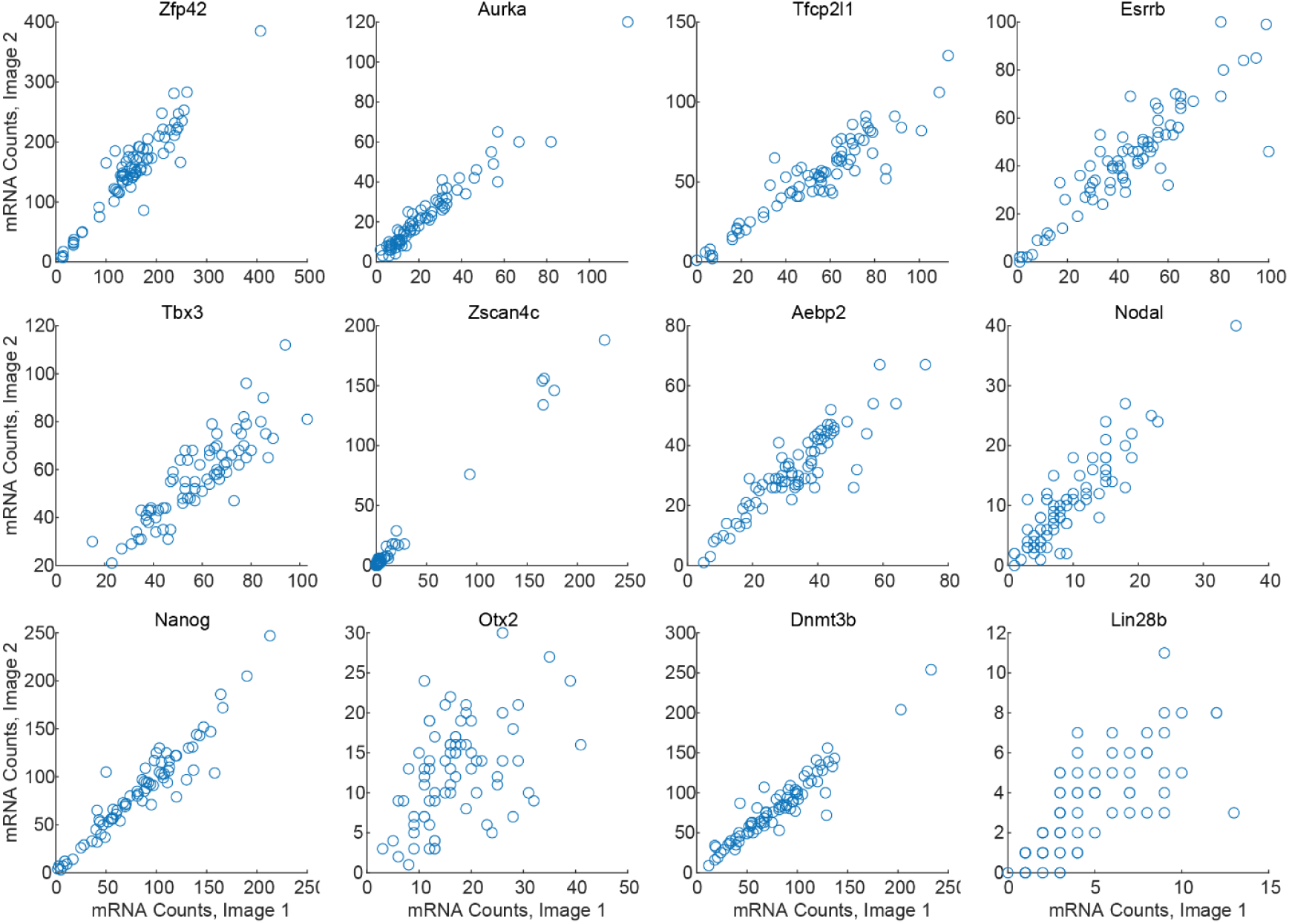
Gene detection is consistent across images. Gene counts as quantified from FISH images by the bigFISH package^59^ are correlated for cells that were measured in multiple images.

